# Switching Functional DNA-Binding Modes by Tuning Protein Order-Disorder Equilibria

**DOI:** 10.64898/2026.08.11.744313

**Authors:** Shilpi Laha, Harshavardhan Madhan, Yuji Itoh, David de Sancho, Satoshi Takahashi, Athi N. Naganathan

## Abstract

Physical remodeling of chromatin by non-histone architectural proteins of the High Mobility Group B (HMGB) family is central to eukaryotic transcriptional regulation. Nhp6A, the prototypical single-HMG-box protein from yeast, harbors both ordered and disordered regions enabling it to bind and bend DNA without sequence specificity. Here, we integrate ensemble experiments, single-molecule FRET, statistical mechanical modeling and atomistic simulations to dissect the structural and functional consequences of context-dependent phosphorylation in the ordered domain and its interplay with the intrinsically disordered region in Nhp6A. We find that Nhp6A occupies a narrow thermodynamic window, with a melting temperature close to the growth temperature of its host organism and high unfolding cooperativity, a feature conserved across the HMG-box family. Phosphorylation extents – mimicked by multisite phosphomimetic substitutions at residue positions conserved across fungal taxa – smoothly tuning the conformational equilibria between at least two different substates in the native ensemble, apart from the unfolded state. This intrinsic plasticity enables close packing of Nhp6A on DNA through two degenerate binding modes, accompanied by two distinct DNA bending geometries. DNA rescues a strongly destabilized mutant, T63D, through favorable intermolecular interactions, thus effectively acting as a chaperone driving folding. Our findings thus reveal a conserved sequence-ensemble-dynamics code in Nhp6A wherein not just stability, but also phosphorylation-induced conformational switching, disordered tail dynamics, and DNA binding-bending closely coordinate chromatin accessibility. The combination of marginal stability, large cooperativity and electrostatic frustration emerges as a design principle to encode charge sensitivity into proteins, and may represent a general strategy for multisite post-translational regulation.

## Introduction

Protein phosphorylation is the predominant post-translational modification (PTM) underlying cellular signaling in eukaryotes. By catalyzing the transfer of a phosphate group from ATP to serine, threonine, or tyrosine residues, protein kinases transduce extracellular stimuli and intracellular signals into alterations in protein activity, interaction specificity, subcellular localization, and stability.^1^ The mechanistic consequences of phosphorylation are highly context-dependent. The introduction of a doubly negative phosphate group can disrupt or promote specific intramolecular contacts, alter surface charge distribution, and modulate protein–protein and protein–nucleic acid interactions.^2^ From a thermodynamic perspective, phosphorylation can shift the conformational landscape of a folded protein, redistributing populations between folded, partially folded, and disordered states,^3–9^ or even its oligomerization status.^10–13^ This phenomenon is particularly common in intrinsically disordered proteins (IDPs) and proteins harboring disordered regions, where the associated energy landscape with multiple minima renders them exquisitely sensitive to charge perturbations.^14,15^ Multisite phosphorylation can further act as a switch or a rheostat, enabling bistable or graded responses depending on the arrangement and cooperativity of phosphorylation sites.^16,17^ In the nucleus, phosphorylation of chromatin-associated proteins that control access to DNA modulates nucleosome dynamics, and interfaces with the broader histone code to regulate transcriptional output.^18^

The members of the High Mobility Group (HMG) protein superfamily — non-histone architectural proteins that remodel chromatin without sequence specificity — are particularly susceptible to phosphorylation-dependent regulation.^19–23^ In the double HMG-box HMGB1, PKC-mediated phosphorylation of S46 and S53 within the folded A-box domain significantly reduces DNA-binding affinity and destabilizes the folded domain.^24^ For single-HMG-box proteins such as HMG-D (Drosophila) and ZmHMGB1 (maize), phosphorylation of serine residues within the intrinsically disordered acidic tail by casein kinase 2 (CK2) and PKC stabilizes an auto-inhibited conformation in which the DNA-binding surface is occluded, revealing that phosphorylation can act as a structural rheostat — toggling between DNA-binding-competent and auto-inhibited states.^25^

Nhp6A, the prototypical single-HMG-box protein in *Saccharomyces cerevisiae* and one of the most abundant nuclear proteins in yeast, binds and sharply bends duplex DNA in a non-sequence-specific manner through minor groove interactions.^26–28^ These interactions are mediated through the ordered region that has a classic three-helix bundle topology with an L-shaped fold and a basic N-terminal intrinsically disordered region (N-terminal IDR; Figure 1A, 1B).^29,30^ This long basic IDR, which results in strong local electrostatic repulsion (Figure 1C), and the lack of a C-terminal extension make Nhp6A differ from virtually all other members of the single-HMG-box protein subfamily. Nhp6A deletion mutants lead to growth defects in yeast^31^ as this single-gene product controls RNA polymerase II-dependent transcription, promotes assembly of preinitiation complexes, and cooperates with the FACT complex to reorganize nucleosomes.^32^ Despite its critical nuclear functions, the structural and functional consequences of phosphorylation in the ordered domain of Nhp6A are unexplored.

This coexistence of ordered and disordered elements in Nhp6A places it at the boundary between structured and unstructured proteins. This makes Nhp6A a unique model system for interrogating how order and disorder cooperate to encode functional versatility, i.e. the non-specific DNA binding and bending. Specifically, is the conformational landscape of Nhp6A poised at the threshold of disorder to exhibit maximal mutational sensitivity? Phosphoproteomic studies in yeast report elevated phosphorylation of Nhp6A under mutational stress, with S26, S41, and T63 identified as the primary modification sites.^33,34^ Notably, T63 in Nhp6A is the structural equivalent of S53 in HMGB1, a residue phosphorylated by PKC.^24^ A key characteristic of these sites is that they are all located within the folded domain: S26 serves as a N-cap for helix 1, S41 is located at the C-terminus of helix 1, and T63 is located in the short loop connecting helices 2 and 3 (see Figure 1A, 1B). Whether these phosphomimetic substitutions perturb thermodynamic stability, DNA-binding affinity, or both remains to be determined. It is likewise unresolved whether any resulting conformational switching behaves as a bistable toggle or a continuous rheostat. Given Nhp6A’s role as an architectural protein, it will also be important to establish whether these mutations subtly alter DNA binding and bending modes. Beyond addressing these specific questions, we develop experimental and computational frameworks that are broadly applicable to interrogating the coupled processes of folding, binding, and bending in non-specific DNA-binding proteins.

**Figure 1.**
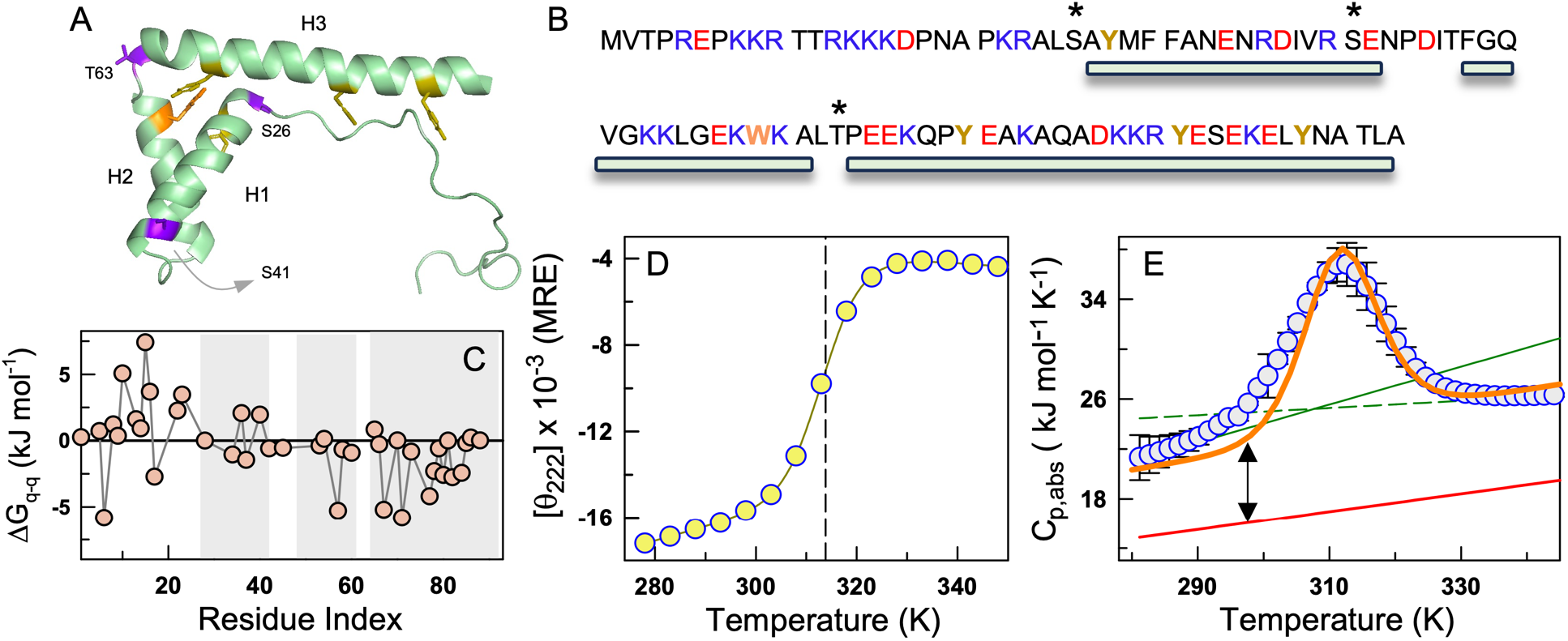
Sequence-structure-thermodynamic features of Nhp6A. (A) Cartoon of Nhp6A with the three helices depicted as H1, H2 and H3 (PDB id:1LWM). The tryptophan and tyrosine residues of interest are highlighted in orange and dark yellow, respectively. The positions mutated in this study are highlighted in purple. (B) Sequence with the acidic residues in red and basic residues in blue. The starred residues – S26, S41 and T63 - are the phosphorylation sites studied in this work. The green bars below the sequence represent the helical regions. (C) Tanford-Kirkwood (TK) electrostatic interaction free energies as a function of protein sequence. Shaded areas represent the boundaries of the helices. Positive values highlight strong unfavorable electrostatic interactions. (D) Thermal unfolding curve monitored by far-UV CD at 222 nm (circles) and the associated fit from a two-state model (curve). The signals are reported in mean residue ellipticity (MRE) units of deg. cm^2^ dmol^-1^. The vertical dashed line represent the melting temperature (*T_m_*) of 313.4 K obtained from a two-state fit. (E) Absolute heat capacity of Nhp6A as a function of temperature (blue circles). The green solid and dashed lines are folded and unfolded baselines, respectively, from a two-state fit. The Freire baseline is shown in red. The double arrow indicates the large enthalpic fluctuations associated with Nhp6A at 298 K. The orange curve is the fit to the bWSME model.

## Methods

### Over-expression and purification of Nhp6A

The gene corresponding to Nhp6A (Uniprot id P11632) from *Saccharomyces cerevisiae*, with sequence MVTPREPKKRTTRKKKDPNAPKRALSAYMFFANENRDIVRSENPDITFGQVGKKLGEKWKALTPEEKQPYEAKAQADKKRYESEKELYNATLA was a gift from Reid C Johnson (UCLA, USA). It was cloned into the NdeI and SpeI sites of the pTXB1 vector (IMPACT™ from New England Biolabs) with a C-terminal intein tag (∼28 kDa). The plasmid containing the gene of interest was transformed into *E.coli* BL21(DE3) cells. A single bacterial colony carrying the transformed plasmid was incubated in 2 L of Luria-Bertani (LB) media containing 50 µg/mL ampicillin and grown at 37°C and 180 rpm in an orbital shaker. The culture was induced with 1 mM IPTG at an OD of 0.8-0.9 at 600 nm. After a 16-hour incubation period at 16°C, the cells were harvested by centrifugation at 8000 rpm for 5 minutes at 4°C. The supernatant was discarded, and the pellet was stored at -80°C. The pellets corresponding to 1 L of culture were dissolved in 100 mL of lysis buffer (20 mM Tris, 1 M NaCl, 1 mM EDTA) with 1 mM PMSF, sonicated and then centrifuged at 11000 rpm for 60 min at 4°C. The lysate was treated with 0.3% polyethyleneimine (PEI) and centrifuged to remove DNA. The clarified lysate was loaded manually (flow rate - 0.8 mL/min) onto a 5 mL chitin resin column previously equilibrated with 20 column volume (CV) of lysis buffer. It was then washed with 5 CV of wash buffer (20 mM Tris, 2 M NaCl), followed by 20 CV of Buffer A (20 mM Tris, 100 mM NaCl). The intein tag was cleaved by incubating the chitin column for 16 hours in 100 mM β-mercaptoethanol dissolved in Buffer A. After the incubation period, the protein was eluted with Buffer A, and the protein fractions were again passed through fresh chitin columns pre-equilibrated with buffer A to remove the free intein tag. The fractions containing the protein were pooled, lyophilized, and loaded onto a Superdex 75 pg size-exclusion chromatography (SEC) column, pre-equilibrated with 150 mM ammonium acetate, pH 8, eluted at a flow rate of 2 mL/min, and the purity of the protein samples were confirmed with a 16.5% tricine SDS gel. The eluted fractions were pooled, lyophilized, and stored in -80°C for further experiments. Mutants were generated by site-directed mutagenesis using primers designed with the NEB base-changer tool. The truncated Nhp6A variant (trNhp6A, where the N-terminal IDR from residues 2-23 is deleted) and phosphomimetic mutants of Nhp6A were purified following the same protocol as the wild type.

All the experiments were performed with filtered and degassed 150 mM ionic strength (IS) buffer (20 mM sodium phosphate with 107 mM NaCl). The lyophilized protein was dissolved in the buffer and filtered through a 0.22 μm syringe filter. The protein concentration was measured using a UV–visible spectrophotometer (Jasco, Inc.) with an extinction coefficient of 11460 M^−1^ cm^−1^ at 280 nm.

### Circular dichroism (CD) and fluorescence experiments

Far-UV CD spectra were recorded in a Jasco J-815 spectropolarimeter equipped with a Peltier system at 5 K temperature intervals from 278 to 368 K in a 1 mm pathlength quartz cuvette at [protein] of ∼12 µM. The protein samples were equilibrated for 2 minutes before spectral acquisition at every temperature. The data for the WT and mutants were fit to a global two-state model with a single folded baseline to estimate melting temperatures (*T_m_*). Fluorescence emission spectra of WT and mutants were acquired in a Chirascan Plus qCD instrument (Applied Photophysics Ltd., UK) equipped with a Peltier system, using a 10 mm pathlength cuvette at a [protein] of ∼10 µM. The protein samples were excited at 274 or 295 nm, with the latter monitoring mostly the tryptophan emission.

### Differential scanning calorimetry (DSC)

Apparent heat capacity profiles were measured at protein concentrations of 64.1 µM and 97.4 µM for the WT in an automated VP-DSC microcalorimeter (Malvern 187 MicroCal VP, NL) at a scan rate of 1.5 K/min. Before the experiment, the samples were buffer-exchanged using a 26/10 HiPrep column (Cytiva) to eliminate residual salts, and degassed for 5 minutes. Buffer scans were acquired before and after every protein scan to ensure that there is no baseline drift. Absolute heat capacity profiles were estimated as documented before.^35^

### Fluorescence anisotropy

Anisotropy experiments were recorded in a Chirascan Plus qCD instrument (Applied Photophysics Ltd., UK) equipped with a polarization detector. 15 bp double-stranded DNA (5’-GGGGTGATTGTTCAG-3’) at a concentration of 40 nM, labeled with Cy3 at the 5’-end was titrated with increasing protein concentrations (1 pM – 100 µM) in a 10 mm pathlength quartz cuvette. This was followed by gentle stirring of the samples for 5 minutes to ensure homogeneity prior to data acquisition at 25 °C. The excitation wavelength was set at 555 nm with a spectral bandwidth of 12 nm, and the emission was recorded at 600 nm.

### smFRET experiments

Single molecule FRET measurements were performed with a home-built confocal system coupled with the Alternating Laser Excitation (ALEX) system.^36^ A pair of laser sources was employed: 488 nm (for donor excitation; OBIS 488 nm LX 100 mW, 1236444, Coherent) and 642 nm (for acceptor excitation; OBIS 640 nm LX 75 mW, 1236445, Coherent). A water immersion objective (NA 1.2, CFI PlanApo 60XC WI, Nikon) was employed to focus the reflected excitation beam inside the 200 µL sample mounted on a glass dish (IWAKI). The fluorescence signal was collected using the same objective and transmitted through a dual-bandpass filter (ZET488/640m, Chroma Technology). The signal was separated into donor and acceptor photons by a dichroic mirror (cut-on wavelength of 593 nm, #67-083, Edmund Optics), and the photons were routed through their respective bandpass filters (donor: FBH520-40, Thorlabs; acceptor: FF01-676/29, Semrock). Two single-photon avalanche diodes (SPADs; SPCM AQRH-14-FC, Excelitas) were used for detection. The SPADs captured individual photons, and their arrival times were recorded by a counter, operated by a homemade LabVIEW program (National Instruments), and stored in the photon-HDF5 file format.^37^ The laser power was adjusted to ∼40 μW and ∼20 μW for excitation of the donor and acceptor fluorophores, respectively.

The lyophilized protein was dissolved in the experimental buffer, and the experiment was performed over a range of concentrations (1 nM - 10 µM) with 15 bp labeled DNA (50 pM) at ∼22°C. The two DNA strands, labeled with Alexa 488 and Alexa 647, were purchased and annealed by heating at 95°C for 5 min, followed by gradual cooling to room temperature for the experiment. To prevent nonspecific adsorption, the IWAKI glass dish was coated with a 0.5% solution of 2-methacryloyloxyethylphosphorylcholine (MPC) polymer (Lipidure CM5206, NOF) in ethanol. The coated dish was then dried by flowing compressed gas over it. A 200 µL sample mixture containing DNA and protein was prepared and loaded into the center of the glass dish. The data acquisition period was set at 30 minutes to collect a sufficient number of bursts.

The data file in the HDF5 format was analyzed with the FRETBursts software.^38^ The local background rate for three different types of photons (donor photons with donor excitation, acceptor photons with donor excitation, and acceptor photons with acceptor excitation) were estimated in windows of 30 s. The burst search was performed for the three channels, provided that the local count rate for 10 consecutive photons exceeded six times the local background rate. For each selected burst, the number of photons in the three channels was estimated after subtracting the local background rates: the number of the donor photons during the donor excitation 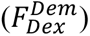, the number of the acceptor photons during the donor excitation 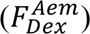 and the number of the acceptor photons during the acceptor excitation 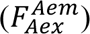 . To exclude the acceptor-only bursts, the first selection was performed by setting a burst size threshold of 20 for 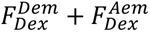. To exclude the donor-only bursts, the second selection was performed by setting a threshold of 20 for 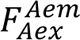. The photon counts for the busts thus selected were used for the calculation of apparent FRET efficiency (*E*) and stoichiometry (*S*)

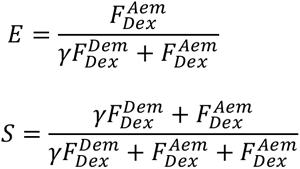

The *γ* value accounts for the differences in quantum yield and detection efficiency between the donor and the acceptor dyes, and was set to 0.48 for this particular experiment.^36^ Note that the *S* values thus estimated depend on the laser intensities used for the donor and acceptor excitation.

### Modeling non-specific protein-DNA binding

The model described below draws upon the theoretical framework of established statistical mechanical models developed earlier to quantify protein–DNA binding equilibria.^39–42^ Assume a DNA fragment of *n* base pairs, and a protein that binds to *ρ* consecutive base pairs on the DNA, in either a minor (denoted as *m*, with a binding free energy Δ*G_m_*) or a major mode (denoted as *M*, with a binding free energy of Δ*G_M_*). In this definition, the minor mode is defined as the weaker binding mode, i.e. Δ*G_m_* < Δ*G_M_*, given the finite size of the DNA fragment. If unbound base pairs on the DNA are represented by the binary variable 0, the microstates of the DNA can be represented by a vector of length *n*, where, each element is 0, *m*, or *M* leading to a total of 3*n* possible microstates. However, given that *ρ* > 1 (i.e. protein binding occludes more than 1 bp) not all 3*n* microstates are possible. We thus combinatorially generate valid microstates descriptions given a value of *n* and *p*, with Ω denoting degeneracy and *O* representing the occupancy. For the simple case of a 5-bp DNA (*n* = 5) binding to a single protein that occludes two sites (*ρ* = 2), the occupancy will be one (*O* = 1) but the multiplicity will be 4 (Ω = 4) as represented by the strings: *mm*000, 0*mm*00, 00*mm*0, 000*mm*. Similarly, it is possible that no protein binds, or one binds in the major mode, two proteins in the minor mode alone or both in the minor and major mode. These effectively represent the macrostate descriptions of the system with the associated multiplicities. See Figure 3D for a pictorial depiction of the same. For the current experimental setup which employs a 15-bp DNA, it is assumed that only a single protein can bind DNA in the major mode as gleaned from the experimental anisotropy data.

The partition function of the system can therefore be written as

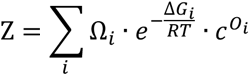

where the summation runs over all the macrostates, and *c* is the protein concentration. The probability of a given macrostate is

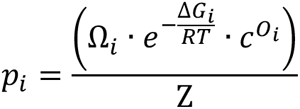

which can be used to calculate the average number of proteins bound

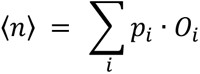

The anisotropy data *S* can be predicted from

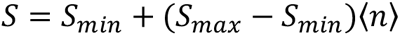

where *S_min_* and *S_max_* represent mean the signal from unbound and maximally bound DNA, respectively. The mean of the anisotropy data at the five lowest concentrations is used to estimate *S_min_* which is fixed for every experiment, including *ρ* that is set at a value of 7 (i.e. occlusion of 7 basepairs on DNA by a single protein estimated from the PDB file 1J5N). The floating parameters are thus Δ*G_m_*, Δ*G_M_* and *S_max_*.

### Block Wako-Saitô-Muñoz-Eaton (bWSME) model

The block WSME model and the associated parameterization are described in detail in recent works.^43,44^ We provide a brief description here. In the original versions of the WSME model,^45,46^ a residue is the elementary folding unit that can either be folded or unfolded and represented by the binary variables *1* and *0*, respectively. This leads to 2^N^ microstates for a N-residue protein, with every microstate represented as strings of *1*s and *0*s. In the more recent versions, we have reduced the phase space accessible to the protein chain by considering consecutive stretches of residues to act as folding units (blocks) and assuming a fixed ensemble defined by microstates that fall under single-sequence approximation (SSA; microstates with single island of folded blocks), double-sequence approximation (DSA; microstates with two islands of folded blocks separated by zeros or unfolded blocks) and DSA with interactions allowed across folded islands (DSAw/L;^47^ same as DSA but including long-range interactions between blocks even if the intervening blocks are unfolded). The stabilization free energy of every microstate ( Δ*G^stab^*) includes contributions from van der Waals interactions (*E_vd_*_W_), all-to-all Debye-Hückel electrostatics (*E_elec_*) and implicit solvation (Δ*G_solv_*). The destabilizing term in the free energy function is entropic (−*T*Δ*S^conf^*) arising from the large penalty incurred in fixing a residue from the unfolded state (*0*) to a folded-like conformation (*1*). These energy-entropy functions are used to calculate the free energy (Δ*G* = Δ*G^stab^* − *T*Δ*S^conf^*) and hence the statistical weight ( *w* = *e*^−Δ*G*⁄*RT*^) of every microstate, and finally the total partition function as *Z* = ∑ *w* where the summation runs over all microstates. Derivatives of the partition function and partial partition functions in turn provide information on residue (block) folding probability, free energy profiles and heat capacity.

In the current work, a block length of 2 is considered resulting in 317,790 microstates for the 93-residue protein. The structure from the PDB id 1LWM is fed into the model and the parameters iteratively modulated to reproduce the experimental heat capacity profile. The final parameters are: van der Waals interaction energy of -143.9 J mol^-1^ per native contact identified with a 5 Å heavy-atom cut-off, entropic penalty of -30 J mol^-1^ K^-1^ per residue, and a heat capacity change of -0.8 J mol^-1^ K^-1^ per native contact (to model implicit solvation). All glycine residues and those identified as non-helical by STRIDE^48^ are assigned an additional entropic penalty of -6.06 J mol^-1^ K^-1^ per residue, while all proline residues are assigned an entropic penalty of zero owing to their limited flexibility. Note that an exact fit was not attempted given the broad heat capacity profile. To simulate the unfolding curves of the mutants, the following van der Waals interaction energies are used while fixing every other parameter to that of the WT: -143.3, -143.0, -141.1 for S26D, T63D and the TM, respectively. To mimic DNA-bound (holo) form of Nhp6A, the entropic penalty is reduced to -28 J mol^-1^ K^-1^ (i.e. lower penalty compared to the apo form) per residue and all other parameters are kept constant. For simulating the free energy profiles of orthologs, only the vdW interaction energy was manually adjusted to match the stability profile of Nhp6A at 298 K while keeping all other parameters constant. The vdW interaction energy per native contact (in J mol^-1^) are provided here in the same order as ortholog ordering in Figure 8F: -146, -146, -144, -147, -143, -143, -146, -145, -143, -146.

### Generating 3D Ensembles from the bWSME model

To identify the structural features of N and N* we resorted to the *Hashi* workflow^49^ that generates 3D ensembles by integrating bWSME model outputs with the RANCH module of ATSAS.^50^ The *macrostate_pools* and *no_of_microstates* define range of reaction coordinate values from which the microstates are chosen from and the number of microstates, respectively. Since more than one structural realization is possible for every microstate (as the *0*s can take up any representation in the 3D space), the parameter *no_of_conformations* is also defined. The final parameters are: *macrostate_pools = [22 27; 28 33], no_of_microstates = 7, no_of_conformations = 3*.

### MD Simulations

We have run atomistic simulations of Nhp6A WT and several of its phosphomimetic mutants (S26D, T63D, and TM, comprising S26D, S41D, T63D) and their complexes with the short 15 DNA duplex described above. Additionally, we have also run simulations of the free DNA duplex and the complex of two copies of the WT protein bound to the DNA. All the simulations were initialized from Alphafold3 predictions^51^ for the protein:DNA complex, except for cases where in the predicted model the protein and DNA were unbound. We used the Amber parambsc1 force field for the nucleic acids^52^ and the Amber14SB for the protein,^53^ together with the TIP3P water model.^54^ Before topology generation the structures were standardised removing the 5’-terminal phosphate of each DNA strand.

All of the systems were fully solvated in a truncated-octahedral box with minimum solute-wall distances of 1 nm for the protein DNA duplex, and neutralized with K^+^ and Cl^-^ up to an ion concentration of 0.15 M. Electrostatics were treated with the Particle-Mesh Ewald method^55^ while Lennard-Jones interactions used a plain 0.9 nm cut-off with no long-range dispersion correction, under the Verlet cut-off scheme. All bonds to hydrogen were constrained with LINCS,^56^ allowing a 2 fs integration time-step. The molecular systems were then simulated using a similar equilibration procedure, including energy minimization, and short simulations (100 ps), first in the NVT ensemble using the velocity-rescale thermostat and then in the NPT ensemble using the Berendsen barostat. Production runs were performed in the NPT ensemble at 298 K using a velocity-rescale thermostat^57^ with a friction constant of 1 ps^-1^ and the Parrinello-Rahman barostat.^58^ All the simulations were run using Gromacs 2024^59^ patched with PLUMED (version 2.9).^60^

For the WT protein, DNA and protein:DNA complex, we have run three replicates of short equilibrium simulations (250 ns) to monitor fluctuations around the AF3 predicted initial states (Table S1). Additionally, conformational sampling was enhanced using two-dimensional well-tempered metadynamics with file-based multiple walkers whose hills files were shared periodically to construct a single history-dependent potential of mean force.^61^ For the apo form of Nhp6A, we used six walkers to sample the fraction of native contacts (*Q*_core_) for C^α^-C^α^ contacts of the folded core (residues 21–93) within 0.65 nm and the coordination between the disordered tail and the folded core. For the protein:DNA complex, we bias the tail-DNA coordination, defined as a continuous coordination number between the N-terminal disordered tail (C^α^+C^β^ of residues 1–20) and the DNA backbone (P+C1’ of all nucleotides on both strands) and the distance between the centres of mass of the two DNA ends (*d*_e2e_) as a proxy for DNA bending. Gaussian hills (height 2.0 kJ/mol, widths σ = 3.0 and 0.25 nm for the two CVs respectively) were deposited every 500 steps (1 ps) with a bias factor of 5 at 298 K, on a grid spanning 0-150 for the coordination and 0-8 nm for *d*_e2e_. Walker bias files were synchronised every 500 steps. To preserve B-form duplex integrity throughout the biased protein:DNA runs, an repulsive wall with an exponential functional form was applied to a DNA inter-strand fraction of native contacts,^62^ using a reference value of *Q*_DNA_=0.9. We have followed the same procedure to simulate the bound state of the S26D and T63D mutants and also for the triple mutant including S41D.Additionally, we run reference multi-walker metadynamics simulations for DNA only using a single CV (*d*_e2e_).

### Sequence analysis

Sequence similarity searches were performed using BLASTp^63^ with Nhp6A as the query against the ClusteredNR database (default parameters), retrieving 100 fungal sequences (taxid: 4751). Sequences were filtered to retain only those containing a single HMG domain and an N-terminal tail. Multiple sequence alignment was carried out using ClustalW,^64^ and sequence conservation was visualized using WebLogo.^65^ Per-residue absolute Shannon entropy was calculated using the Shannon Entropy-One web server (https://www.hiv.lanl.gov/content/sequence/ENTROPY/entropy_one.html) to quantify positional variability across the alignment. Ten homologs were manually selected, and their structures were predicted using the AlphaFold 3 server.^51^ Electrostatic interaction free energies for each orthologs were then computed via Tanford–Kirkwood calculations^66^ with the predicted structure as input with the following parameters: pH 7.0, 298 K, ionic strength of 150 mM.

### Simulation of stability-cooperativity scenarios

For a model WT system of 67 residues, we fixed the heat capacity change at 40 J mol^-1^ K^-1^ per residue and considered two melting temperatures – 333 K or 313 K (high and low *T_m_*, respectively) - and unfolding cooperativity values of 2.9 and 1.45 kJ mol^-1^ per residue (high and low Δ*H_m_*, respectively). We then simulated changes in stability as a function of temperature using the standard Gibbs-Helmholtz relation and derived folded state probabilities for the different scenarios. Each successive mutant in every scenario is destabilized by 4 K in *T_m_* such that the *T_m_* of the n^th^ mutant is *T_m_*(*mut*) = *T_m_* − 4*n*, while fixing all other paramters. For the scenario where the Δ*H_m_* is assumed to be dependent on the degree of destabilization we use Δ*H_m_*(*mut*) = (1 − 0.1*n*)Δ*H_m_*.

## Results

### The native ensemble of Nhp6A

To characterize the conformational landscape of WT Nhp6A, we first determine its thermodynamic stability from far-UV CD thermal melts. This yields a melting temperature (*T_m_*) of 313.4 ± 0.5 K from a two-state model fit and a thermodynamic stability of 6.5 kJ mol^-1^ at 298 K, suggestive of a marginally stable system (Figure 1D). The unfolding enthalpy at the midpoint (Δ*H_m_*) is estimated to be 186 ± 14 kJ mol^-1^, which translates to ∼2.8 kJ mol^-1^ per residue for the 67-residue structured part of the protein (i.e. excluding the disordered N-terminal tail). This number is comparable to the expected 2.9 kJ mol^-1^ per residue from size-scaling arguments,^67^ indicating that the folded core of Nhp6A has a packing density comparable to that of larger proteins. Consistent with this, the fluorescence emission maximum of the single tryptophan residue, W59, is centered at 330 nm and remains unchanged over the temperature range of 278– 300 K (Figure S1). A large excess heat capacity peak is hence visible in absolute heat capacity (*C_p_*) measurements (Figure 1E) with an amplitude comparable to that of Ubiquitin at pH 3.0.^68^ The baselines cross when fitting the absolute heat capacity to a two-state model; this could be a consequence of non-two-state nature of the system or from the large enthalpic fluctuations (*σ_H_*) of the N-terminal tail and the third helix that it contacts with. We observe a ∼8 kJ mol^-1^ K^-1^ difference between the heat capacity expected from a well-folded protein (Freire baseline) and the experimental data at 298 K. From the relation 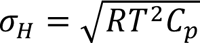, the difference translates to ∼25% more enthalpic fluctuations in the native ensemble of Nhp6A relative to a well-folded protein.

To further probe the structural integrity of the protein, we predict the heat capacity profile through the block Wako-Saitô-Muñoz-Eaton (WSME) model with an ensemble of 317,790 microstates derived from the native structure (see Methods).^43,44^ The broad thermogram precluded an exact agreement with the model predictions that is substantially sharper (orange curve in Figure 1E). Consequently, the conformational heterogeneity inferred from the model represents a lower-bound estimate. The free energy profile at 298 K, generated by projecting the microstates on to the reaction coordinate, the number of structured blocks, shows a broad native ensemble with two substates – one distribution at ∼23 structured blocks (N) and another at ∼31 structured blocks (N*; Figure 2A, 2B). The barrier height for unfolding is large (20 kJ mol^-1^) in agreement with the sharp thermogram observed.

**Figure 2.**
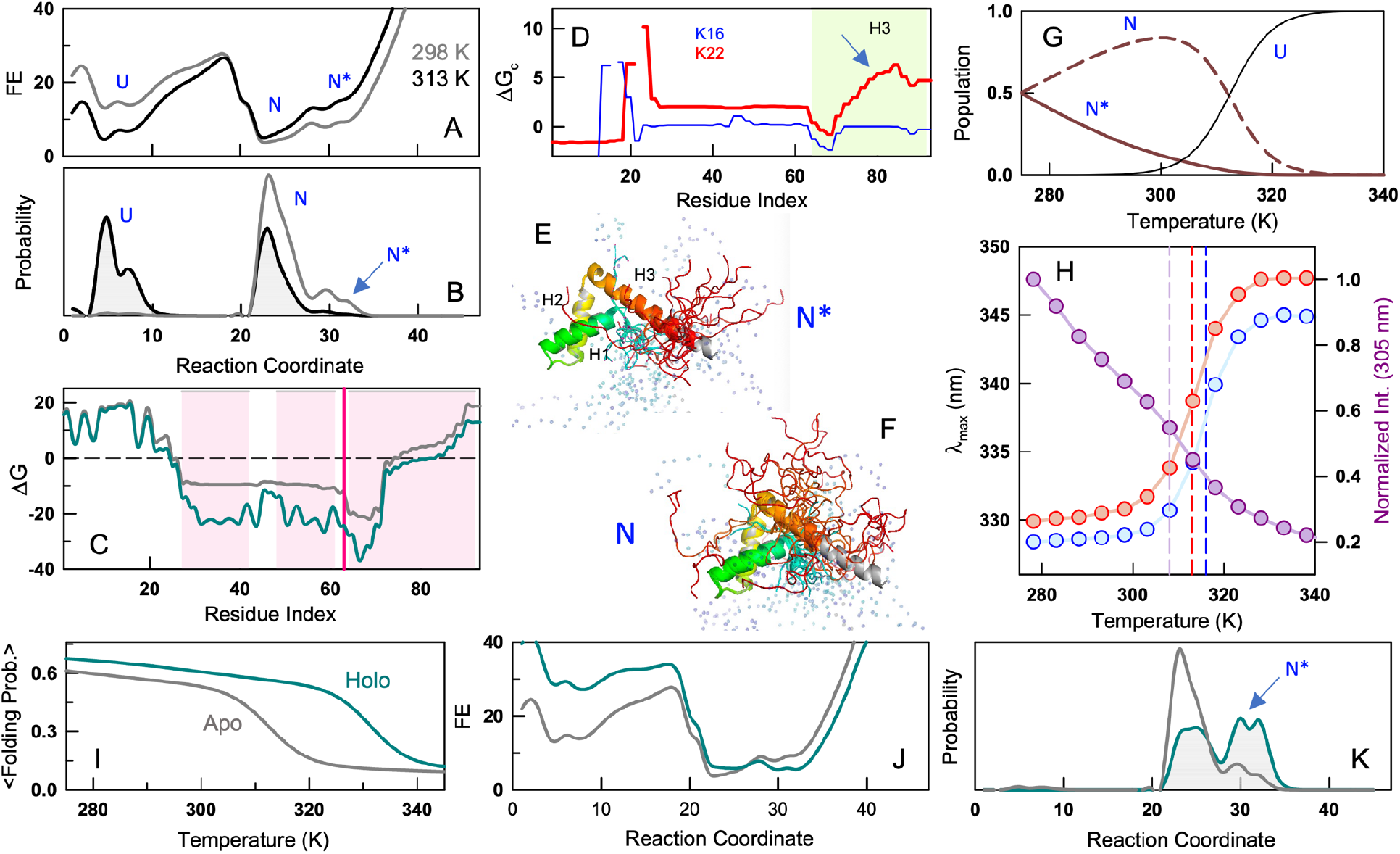
The Nhp6A native ensemble and stability determinants. All free energy units are in kJ mol^-1^. (A, B) The bWSME model predicted one-dimensional free energy profiles as a function of the reaction coordinate (RC), the number of structured blocks, at two different temperatures (panel A) and the corresponding probability densities (panel B). Note the population of an additional macrostate at RC values >30 at 298 K (arrow). (C) Residue-level stability profiles for apo (gray) and holo (dark cyan) Nhp6A. The shaded pink areas represent the three helices. The dark pink line denotes the position of the residue T63. (D) The effective coupling free energy between the residue K16 and all other residues (blue). The red curve plots the same for K22 in the ordered segment of IDR, with the arrow highlighting regions in H3 (shaded area) that display strong coupling with K22. (E, F) Predicted structure of the macrostates at 28<RC<33 (N*; panel E) and 22<RC<27 (N; panel F) by combining the 1D strings from the bWSME model with the RANCH output. The native structure is shown in gray. The disordered residues 1-20 in the IDR are shown as spheres. (G) Predicted population of the substates as a function of temperature from the bWSME model. (H) Thermal melts as monitored by emission maxima on excitation at 274 nm or 295 nm (blue and red, left axis), and the fluorescence intensity at 305 nm (purple, right axis). Vertical dashed lines signal the apparent melting temperatures. (I) Mean folding probability as a function of temperature for apo and holo forms of Nhp6A denoted by gray and cyan, respectively. (J, K) Free energy profiles (panel J) and probability densities (panel K) at 298 K for the apo and holo forms following the color code in panel I. Note that the more folded macrostate (RC>30) is more populated in the bound form.

Residue-level analysis of local stability (ΔG) indicates that, on average, the third helix is less stable than the first two helices (Figure 2C), but with one exception. The N-terminal segment of the third helix is predicted to be highly stable, which has its origins in an *i* − *i* + 4 electrostatic interaction between K67 and E71, with both the residues located in an favorable environment with the Tanford-Kirkwood (TK) electrostatic interaction free energies of -5.7 and -5.6 kJ mol^-1^, respectively. We hypothesize that this region acts as an electrostatic staple to stabilize the N-terminal region of the third helix, while allowing for conformational flexibility in the C-terminal half of helix 3 (H3). Additionally, residues 21–26 — part of the N-terminal IDR termed ‘the extended strand’,^30^ immediately abutting the C-terminal end of H3 — span both order (negative ΔG) and disorder (positive ΔG). The shifts in local stability values suggests that the conformational status of H3 could be coupled to the degree of local disorder in this adjoining segment, despite it appearing ordered in the NMR structure 1LWM. To further quantify this, we calculate the effective coupling free energy (Δ*G_c_*)^69^ that accounts the pair-wise folding probabilities of residue pairs across the entire ensemble of 317,790 microstates. As an example, we consider two residues – K16 and K22 – both part of the N-terminal IDR but exhibiting high and low structural variability, respectively, in the NMR models. K16 is strongly coupled to sequentially proximal residues (Δ*G_c_*>0), negatively coupled to residues N-terminal to it and not coupled to all other residues (Δ*G_c_*∼0; blue Figure 2D). K22 is likewise strongly coupled to its sequential neighbors, but additionally displays a distinct and favorable coupling pattern to residues 77-93 in H3 (red in Figure 2D), highlighting long-range coupling between these two structurally distinct elements.

To visualize the ensemble, we predict the structures of N and N* using the RANCH module of ATSAS^50^ with the binary strings from the bWSME model as the input. Specifically, three microstates with the largest statistical weights within the subensembles are chosen and fed into the RANCH module that provides three-dimensional realizations based on regions of the protein that are folded (*1*s) and unfolded (*0*s). It can be seen that the microstates in the subensemble N* have at least 1-2 turns at the C-terminus of the H3 unfolded with the rest of the protein exhibiting a near native architecture (Figure 2E). The sub-ensemble N is characterized by partial unfolding in 3-4 turns at the C-terminal of H3 (Figure 2F), as a consequence of loss of stabilizing interactions with residues 21-26 at the N-terminal IDR that adopt more disordered conformations.

At 298 K, the population of N* is ∼16% that decreases to zero at 313 K, the denaturation midpoint reported by far-UV CD (Figure 2G). The population of N displays a trend wherein it increases with temperature, reaches a maximum at ∼300 K and decreases again (Figure 2F), consistent with the features of a partially structured state. The folded domain of Nhp6A has two distinct substructures – the folded major wing with Y28, W59 and Y70 forming a core (H1 and H2), and the minor wing harboring Y81 and Y88 in H3. In line with this expectation, the fluorescence emission spectrum of Nhp6A at 274 nm (both tyrosine and tryptophan residues are excited) is blue shifted relative to 295 nm excitation (fluorescence is dominated by W59) (Figure S1C). Any change in fluorescence emission maxima and intensity are a weighted average of both the substructures, and differences in melting temperature would report on their relative stabilities. In the range of temperatures between 278 – 300 K where the emission maximum of W59 is temperature-independent (i.e. the major wing is fully folded), the fluorescence emission intensity at 305 nm (dominated by tyrosine residues) decreases by ∼40% with a sigmoidal-like transition and a lower *T_m_* of 308 ± 1 K (from a first-derivative analysis, and compared to 313.4 by far-UV CD; Figure 2H). Even the emission maxima display slightly different melting temperatures with *T_m_* values of 316 and 314.5 K for 274 and 295 nm excitation, respectively (Figure 2H). This suggests that the minor wing tyrosine residues get solvent exposed at lower temperatures due to melting of the C-terminal half of H3 aided by the larger flexibility of the structured part of the N-terminal IDR. The weaker stability of H3 is in agreement with similar interpretations from comparisons involving apo-holo forms of Nhp6A,^30^ fluorescence and NMR experiments on Sox5,^70^ and high-resolution NMR spectroscopy on Sox2.^71^

The observations above support the applicability of the model in describing the conformational landscape of Nhp6A. The binding of Nhp6A to DNA is expected to enhance the rigidity of the entire protein through a combination of strong van der Waals packing and favorable charge-charge interactions between the basic residues in the protein and the phosphate backbone of DNA. Using the WSME model, it is possible to predict the effect of DNA binding on the stability of the protein by mimicking the rigidification (see Methods). The resulting free energy profiles confirm a stabilization of the protein (dark cyan in Figure 2I) and an increase in the population of N*, the more folded conformation (Figure 2J, 2K). Under these conditions, it appears that the native ensemble of Nhp6A populates both N and N* to equal extents hinting at the possibility of two similar modes of DNA-binding.

### DNA binding

We verify the predictions above by performing far-UV CD experiments on Nhp6A in the presence of a 15-bp double-stranded DNA. Spectral changes, which are independent of protein concentration, are apparent when the apo and holo forms of Nhp6A (in 1:1 molar ratio with 15-bp DNA) are monitored by far-UV CD, and the red-shift of holo-Nhp6A CD spectrum is indicative of secondary structure acquisition (Figure 3A). However, the intensity changes at 222 nm are minimal suggesting that there are compensatory effects. Specifically, if Y81 and Y88 in the C-terminal helix acquire structure on binding DNA, they would interfere with secondary structure content estimates.^72^ The melting temperature of Nhp6A, on the other hand, increases by 15 K in the presence of DNA, signaling strong structural stabilization in the holo form (Figure 3B).

The binding of Nhp6A to Cy3-labeled 15-bp DNA is monitored by fluorescence anisotropy. Increasing concentrations of the protein results in a bi-phasic transition with the first transition displaying an inflection point at ∼120 nM and the second at ∼16 µM (Figure 3C). The clear separation in the concentration regimes is evidence that two molecules bind, but with different binding free energies. The trNhp6A, the variant in which the N-terminal IDR (residues 2-23) is deleted, does not bind DNA in the concentration ranges tested, highlighting the critical role of the tail in mediating strong interactions for a stable complex. A simple two-site binding model is not applicable for Nhp6A, as DNA binding is non-specific with no sequence preference. To better quantify the binding free energies, we developed a statistical mechanical model that considers two binding modes – a major binding mode *M* with a binding free energy Δ*G_M_* and a minor binding mode *m* with a binding free energy Δ*G_m_* – that leads to 5 macrostates: no protein bound (macrostate 0), one bound protein (*m* or *M*) and two bound proteins (*mm* or *Mm*) (Figure 3D). Note that the macrostate *MM* corresponding to two proteins binding in the major mode is not considered, as it would not manifest as two different transitions that is experimentally observed. Second, we assume no coupling between the two binding modes, i.e. no cooperativity factor is introduced. The macrostate statistical weights are additionally weighted by their corresponding multiplicities, which account for the different possible arrangements of a non-specifically binding protein on DNA and the number of occluded binding sites (see Methods).

**Figure 3.**
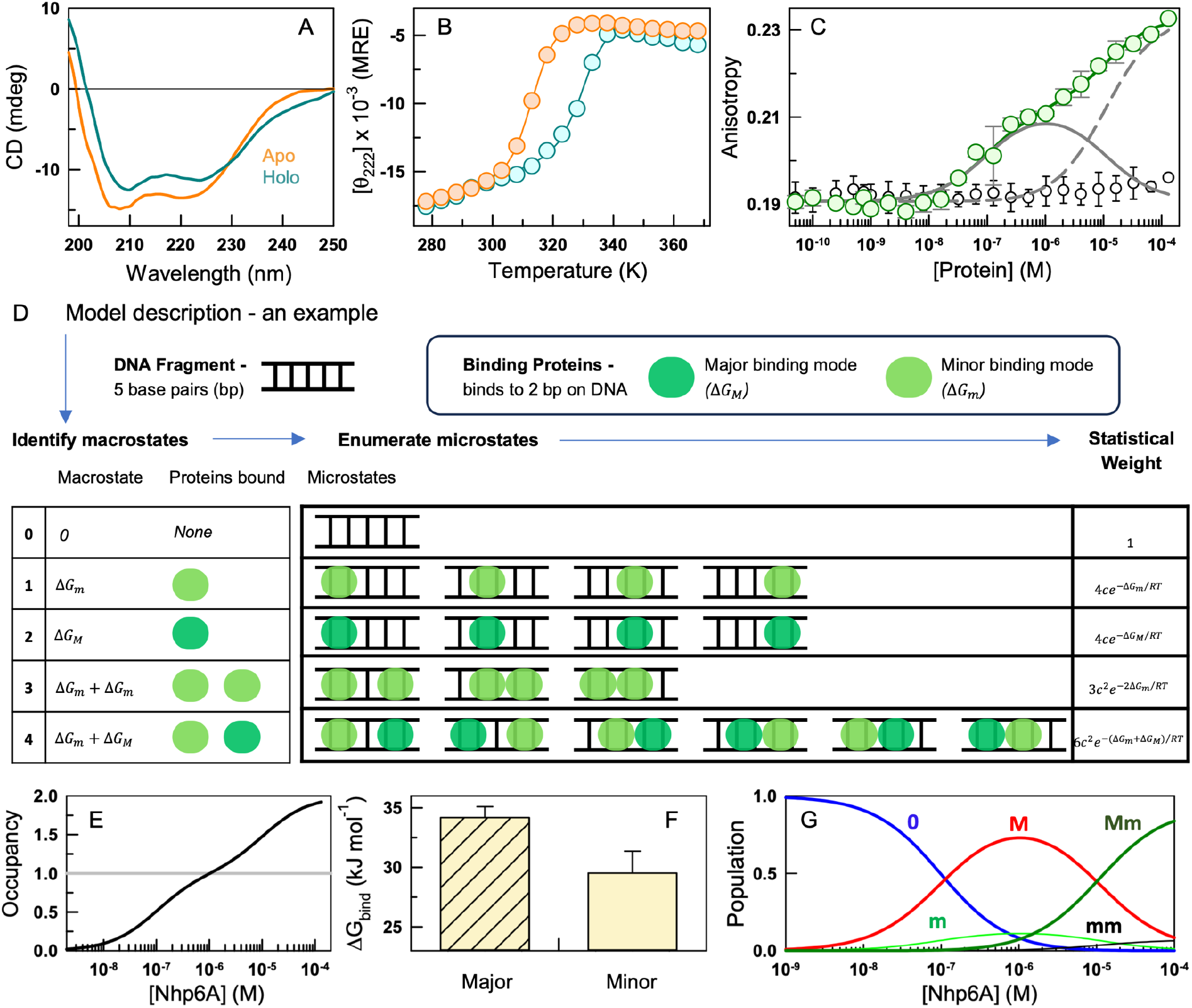
DNA binding modes. (A) Far-UV CD spectra of Nhp6A in the absence (apo, orange) and presence (holo, green) of DNA at 298 K. Note the spectral shift towards a more folded structure in the holo form. (B) Far-UV CD thermal unfolding curves of apo and holo forms of Nhp6A monitored at 222 nm following the color code in panel A and reported in MRE units. (C) Anisotropy of Cy3-labeled 15-bp dsDNA (circles) as a function of Nhp6A (filled circles) and trNhp6A (open circles) concentrations at 298 K. The curve through the points is a fit from the model described in panel D. The solid gray and the dashed lines represent the signal from population of one Nhp6A and two Nhp6A molecules bound to DNA, respectively. (D) Model for non-specific DNA binding considering an exemplary case of a protein with two binding modes – major (*M*) and minor (*m*) each occluding 2 bps - binding to a 5 bp DNA, with the binding free energies ΔG_<_ and ΔG_=_, respectively. Under these conditions, only specific binding ‘states’ are allowed, i.e. the ensemble is fixed, as observed in the table corresponding to the microstates. The free energies and signals are then modulated to reproduce the experimental anisotropy profile (see Methods). The variable *c* in the column to the right is the protein concentration. (E) The average number of protein molecules bound to DNA as a function of protein concentration. (F) Extracted binding free energies of the two binding modes. (G) Macrostate populations as function of Nhp6A concentration.

Within this model framework, the shape of the anisotropy curve is determined by the interplay between the number and nature of (major and minor) binding modes, while the occupancy accounts for the total amplitude. Fitting the anisotropy data to this model results in a very good agreement (dark green curve in Figure 3C) with the occupancy going from near zero at [Nhp6A]<10 nM and reaching up to two at 100 µM (Figure 3E). We estimate the binding free energies to be 34.2 ± 0.9 and 29.5 ± 1.8 kJ mol^-1^ for the major and minor binding modes, respectively (Figure 3F). The predicted macrostate populations display a distinct trend expected of concentration dependent processes (Figure 3G). The population of *M* increases at the expense of ‘no binding’ (0) starting from 1 nM [Nhp6A] (Figure 3G), reaches a maximum at ∼1 µM and decreases at higher concentrations; this decrease is a consequence of the replacement of singly bound species by doubly bound species, with the population of *Mm* dominating over that of *mm*. Similarly, the weaker binding mode with a single bound protein (*m*) is minimally populated in the range of concentrations explored. The second transition in the anisotropy isotherm therefore has its origins in weaker and presumably non-specific binding of Nhp6A molecules. Given that the binding free energies are different by only ∼2*RT*, we expect the structural features of the two binding modes to be similar.

### DNA bending elucidated by smFRET measurements

While ensemble anisotropy experiments can report on Nhp6A binding to DNA, they do not capture any accompanying bending of the duplex. To probe this directly, we labeled DNA with Alexa488 and Alexa647 at the 5′ ends of the two complementary strands, using the dye pair as a molecular ruler in freely diffusing single molecule FRET experiments. Protein binding that alters the relative inter-dye distance would then manifest as a change in FRET efficiency, providing a readout of the extent of DNA bending. The FRET efficiency histogram of the doubly labeled DNA is unimodal with a peak centered at ∼0.12 units (Figure 4A, top panel). On addition of the protein, the mean position of the histogram shifts to the right with significant broadening (Figure 4A, lower panels, and Figure S2A). These features are better seen in the plot comparing the normalized distribution of DNA-alone and DNA+10 µM Nhp6A (Figure 4B). While the former distribution is comparable to the theoretical shot noise width assuming a single component, the latter distribution is broader than the theoretical width, suggesting the necessity of at least two states to explain the latter distribution (Figure S3). Following this, we performed a global fit to the concentration dependent data assuming two- or three-Gaussians with the variance of the distributions fixed to that of the DNA-only histogram and floating the mean positions. The sum of squared residuals (SSR) value is higher for a global two-Gaussian fit than that of a three-Gaussian fit, i.e. *SSR*_2_⁄*SSR*_3_ = 2.16, with a P-value less than 0.0001 from the F-test statistic that accounts for both the number of data points and free-floating parameters (Figure 4D). Thus, a three-Gaussian fit, whilst requiring more parameters, accounts for the experimental distribution better than a two-Gaussian fit.

**Figure 4.**
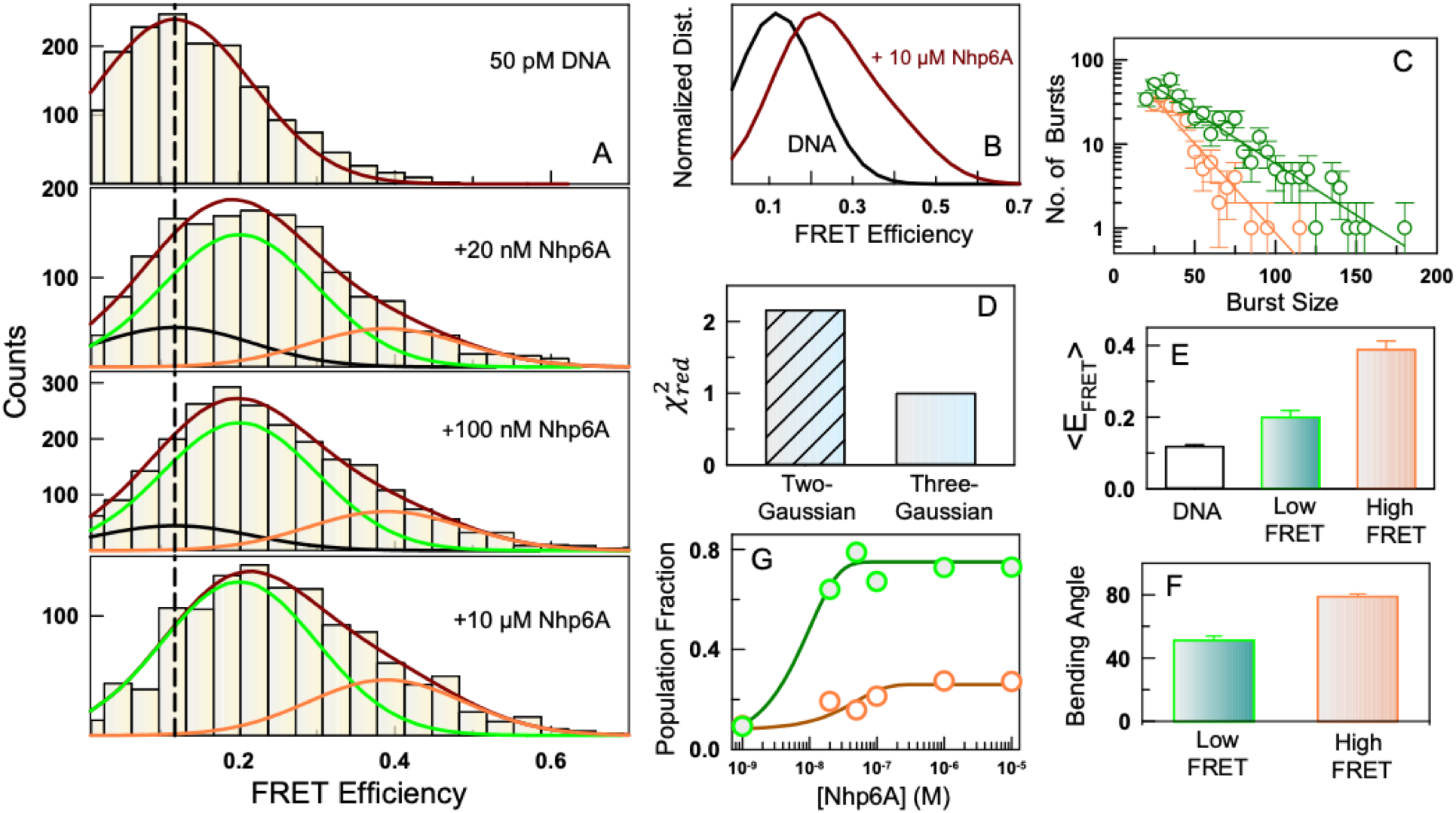
DNA bending from smFRET experiments. (A) Histogram of FRET efficiency distributions for the 15-bp DNA labeled with Alexa488 and Alexa647 at the indicated protein concentrations and at 295 K. The black, green, and orange curves represent Gaussian components corresponding to the DNA and the two different populations with lower and higher FRET efficiencies, respectively. The vertical dashed line corresponds to the mean FRET efficiency of the DNA (i.e. in the absence of the protein). The maroon line is the global fit (see main text for details). (B) Normalized data highlighting the shift in FRET efficiency in the presence of the protein (dark red) compared to the DNA alone sample (black). (C) Burst size distributions of 15-bp DNA in the presence of 10 µM Nhp6A. Detected bursts were categorized into two regions based on with their FRET efficiency *E*: 0.0<*E*<0.20 and *E*>0.40. Bursts with *E* less than 0.20 and larger than 0.4 are shown in green and orange, respectively. Solid lines represent single-exponential fits to the data points. Error bars represent the square root of the number of bursts in each bin. (D) A global three-Gaussian fit provides a better description of the data than a two-Gaussian fit, as indicated by the lower reduced chi-squared value. (E) The mean FRET efficiencies of the three extracted distributions. (F) Bending angles (in degrees) corresponding to the low and high FRET efficiency states. (G) Population fractions of the lower and higher FRET states as a function of protein concentration, following the color code in Panel E. The curves are shown to guide the eye.

To confirm the presence of distinct species for the distribution in the presence of 10 µM Nhp6A, we conducted the burst size analysis for bursts whose efficiencies are less than 0.2 and larger than 0.4 corresponding to the green and orange distributions, respectively (Figure 4C). The slope of the burst size distribution plot (the total number of photons detected during a single burst) on a semi-logarithmic scale is inversely proportional to both the brightness and diffusion time of the sample.^73^ The slope of the bursts with efficiencies below 0.20 (green) is steeper than that of the bursts with efficiencies above 0.40 (orange), showing the distinct difference of the two groups of the bursts. The steeper slope observed for the high-FRET population is most likely due to the quenching of Alexa488 fluorescence (Figure S2B). Thus, the analysis demonstrate that the fitting of the FRET efficiency distribution requires at least three Gaussians, one for 15 bp-DNA alone and additional two corresponding to the DNA–Nhp6A complex.

The resulting mean FRET efficiency values for the three Gaussian-distributions are 0.12, 0.20 and 0.39, respectively, with the latter two accounting for the two additional species identified in the presence of the protein that broaden the histogram (Figure 4E). Using the mean values and the law of cosines,^27^ we estimate the bending angle of DNA to be 52° and 79°, respectively (Figure 4F). The corresponding population fractions are higher for the low FRET state (lower DNA bending) compared to the high FRET state (higher DNA bending; Figure 4G). Our results thus reveal that the binding of Nhp6A to the 15-bp DNA leads to two distinct substates, one which bends the DNA more than the other. The high FRET substate does not arise from the binding of two proteins as this species is observed even at 20 nM concentration. Whilst bending of DNA has been reported for Nhp6A,^27,28^ this is, to the best of our knowledge, the first report where two distinct subspecies are observed along a bending coordinate. The population fractions flatten out beyond 100 nM indicating that the bending event observed in smFRET does not carry information on the number of proteins bound. Overall, smFRET studies reveal that the DNA-bound ensemble is inherently heterogeneous populating not just five different macrostates depending on the concentration, but also resulting in two DNA bending extents.

### Simulations shed light on the heterogeneous binding landscape

To shed light on inferred unstructuring in the apo state and induced DNA bending, we have run atomistic MD simulations of free Nhp6A and its complex with the 15-bp DNA duplex. In both the apo and holo systems, we have run batches of equilibrium simulations and also enhanced the sampling using the metadynamics technique with multiple walkers to explore the slow degrees of freedom involving the IDR at the N-terminus of Nhp6A (see Methods). In the apo form, the native state ensemble predominantly involves fluctuations in the number of IDR-folded core contacts (Figure 5A). Even if we are preventing the system from sampling more partially structured and unfolded states (we used an upper wall that, in practice, does not allow the protein core RMSD to exceed 6 Å), a decrease in the fraction of native contacts (Qcore) is still possible. The decrease in the native contacts is predominantly due to a loss of helical structure primarily in helix H2 (Figure 5B, inset and 5C), which is involved in the binding to DNA in the major mode. These conformational excursions likely represent the earliest structural changes in the apo form and could serve as precursors to the more complex structural transitions observed experimentally. Indeed, NMR CPMG relaxation dispersion experiments in the orthologous Sox2 protein reveal a minor state (∼1% population) with perturbations spanning several residues in H1 and H2,^71^ hinting at conformational heterogeneity even in the folded core of HMG proteins. This could be further aided by the presence of three glycine residues in H2 of Nhp6A, which is unusual for a helix.

Simulations of the complex formed by Nhp6A with DNA indicate that the IDR also has an important role in binding, consistent with experimental observations (Figure 3C). In this case, we find that the DNA duplex indeed bends in presence of Nhp6A relative to reference simulations for the DNA alone (Figure 5D). We note, however, the bending angle observed in simulations is more modest than that derived from experiments, shifting from <10° for free DNA to ∼30° in presence of Nhp6A. The greater stiffness of the DNA in the simulations may be due to the restraining potential we have introduced to avoid the rupture of Watson-Crick base pairing that might result in DNA melting. Still, there is a very clear modulation of bending by the Nhp6A. Importantly, in the two-dimensional free energy landscape we find that bending is possible only upon formation of IDR-DNA contacts (Figure 5E). Positively charged residues in the IDR engage the polyphosphate backbone along one face of the duplex, neutralizing the local charge and driving DNA bending (Figure 5F).^75,76^ These results suggest that the IDR has a decisive role not only in the apo protein’s conformational ensemble but also of the structural ensemble adopted by the partner DNA molecule upon complexation.

**Figure 5.**
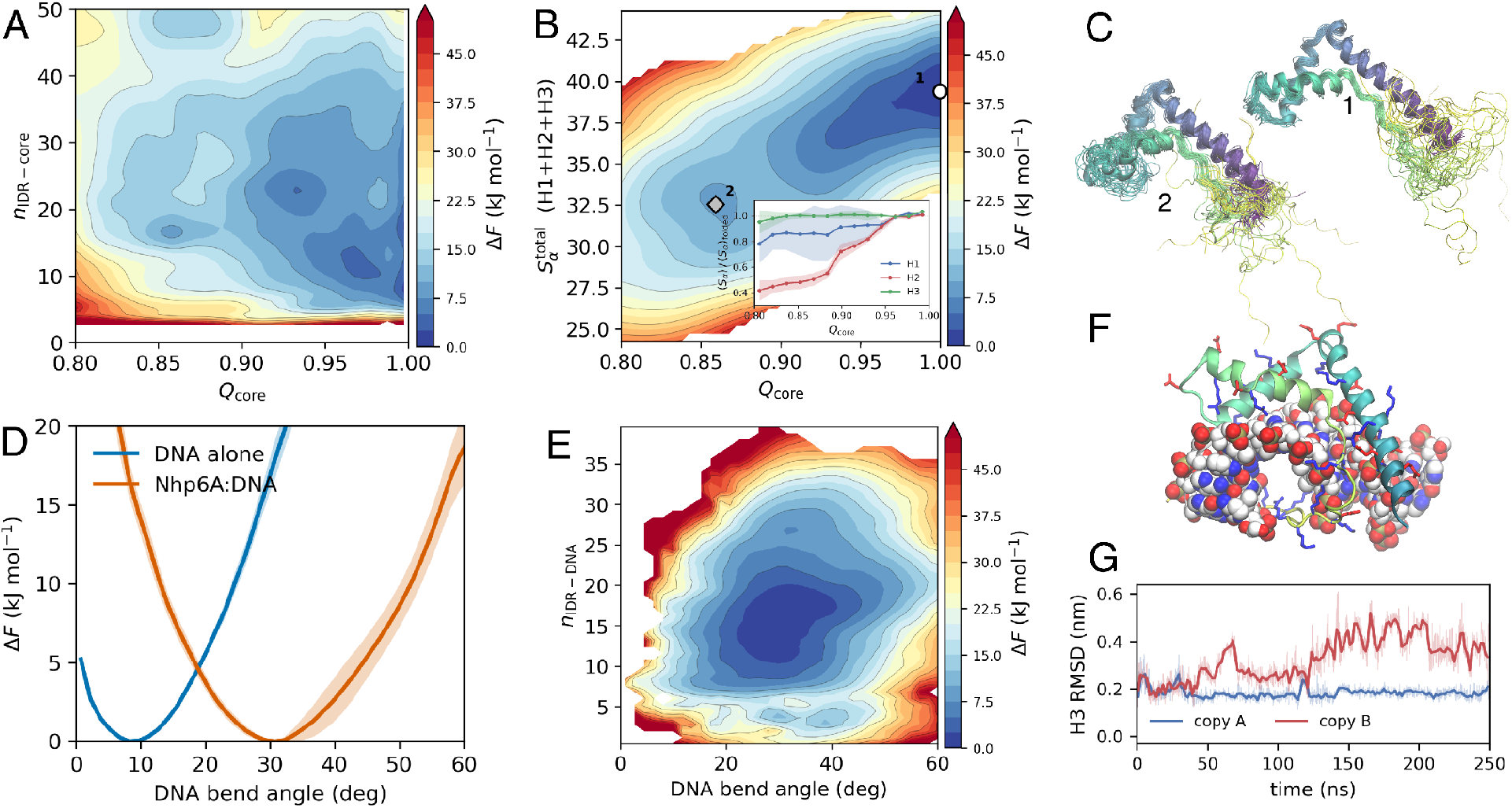
Atomistic simulations of Nhp6A in the apo and holo states. (A-B) Free energy landscape from metadynamics simulations of apo Nhp6A, using the fraction of native contacts of the folded core (*Q_core_*) and the coordination number for core-IDR contacts (*n*_IDR-core_; panel A) and helicity (*S_a_*^total^;^74^ panel B). The inset in panel B is the average helicity relative to that in the AF2 prediction as a function of *S_a_* for the different helices. (C) Medoids and representative snapshots of the dominant state (1) and a metastable state of lower helicity (2). (D) Potential of mean force of the Nhp6A:DNA complex for the projection on the bending angle. (E) Free energy landscape for the protein:DNA complex as a function of the bending angle and the number of IDR-DNA contacts (*n*_IDR-DNA_). (F) A representative snapshot of the DNA:Nhp6A complex highlighting the interaction between the positively charged residues (red and blue sticks) of the IDR (yellow) and the DNA (spheres). (G) RMSD of helix 3, with respect to the starting structure, for the two Nhp6A molecules in the 2:1 protein-DNA complex in one of the short equilibrium trajectories.

To explore the origin of the observed behaviour of the DNA at high protein concentrations, we have run an additional set of equilibrium simulations for the complex formed by DNA with two copies of Nhp6A. We started from the AF3 prediction, where the two protein units are symmetrically bound in the same mode, as a putative model for the complex structure, despite the low pTM and ipTM scores (0.4 and 0.26, respectively). We find that this complex experiences large distortions, particularly in helix H3 (Figure 5G) in timescales of just over a few hundred nanoseconds, likely due to a combination of steric hindrance and positive charge accumulation (each protein copy is +8). This supports the assumption in the statistical mechanical model of DNA binding presented above that the macrostate with two proteins bound in the major-mode makes negligible contributions to the ensemble.

### Rheostat-like modulation of stability by phosphomimetic mutations

Phosphoproteomic studies report that Nhp6A is phosphorylated at three different sites - S26, S41 and T63 (Figure 6A).^33,34^ S26 is located close to the N-terminus of the first helix forming the N-cap and in direct contact with the DNA in the holo form, S41 is at the C-terminus of the first helix and T63 is located at a unique location in the short loop connecting second and third helices. The latter two do not interact with the DNA in the available solved structure 1J5N. Tanford-Kirkwood (TK) electrostatic interaction free energy calculation reveals that the phosphomimetic mutants S26D and S41D have little effect on the native electrostatic interactions (Figure S4), while T63D induces substantial destabilization (positive free energies in Figure 6B). It is interesting to note that T63 is very close to the first two turns of the third helix where a large fraction of the stabilization energy is localized (vertical red line in Figure 2C). It is therefore expected that T63D mutation will destabilize the protein more than the other mutants.

To test for this, we acquired far-UV CD spectra for all of single, double (DM), and the triple (TM) mutants. The intensity of the spectra and the shape vary in a manner consistent with enhanced destabilization as one goes from the WT to the TM (see Table S2 for mutant-specific information including fitting errors). Specifically, the WT exhibits a mean residue ellipticity of -14,900 deg. cm^2^ dmol^-1^ at 303 K while the TM displays a signal which is nearly one-third (-5600 deg. cm^2^ dmol^-1^), and with the spectral minimum shifted to 202 nm from 208 nm (Figure 6C). A global two-state model fit with a shared folded baseline and heat capacity change is able to account for the spread in the melting temperatures, with the *T_m_* ranging from ∼314 K for the WT to ∼294 K for the TM (Figure 6D). Amongst the single mutants it is the T63D which is strongly destabilized with a Δ*G* of just 2.9 kJ mol^-1^ at 298 K, in very good agreement with the TK model expectations (Figure 6E). The Δ*G* of TM is -1.5 kJ mol^-1^ compared to the WT Δ*G* of 6.5 kJ mol^-1^, revealing that the phosphomimetic mutations switch the native ensemble from being fully folded to mostly disordered in a graded manner. Interestingly, the fluorescence *λ_max_* of W59 does not change significantly even at the lowest temperatures (Figure S5) as opposed to the far-UV CD signal at 222 nm that reduces by 4000 deg. cm^2^ dmol^-1^. This observation suggests that the hydrophobic core of the protein is intact even in the most destabilizing variants, and that structural changes are localized in regions farther away from the core.

The standard deviation of the folding probability across the variants as a function of temperature peaks in the temperature range 303−308 K (Figure 6F), which is very close to the optimal growth temperature of *S. cerevisiae* (∼303 K). The correspondence between the optimal growth temperature and maximal tunability, though possibly coincidental, is one an organism could exploit for functional or regulatory advantage. Nhp6A in its native form rarely samples the unfolded ensemble given the large thermodynamic barrier separating the two ensembles (Figure 6G). However, upon phosphorylation or introducing phosphomimetic mutations, the populations tend towards a more unfolded states (U) (Figure 6G, 6H). The protein, depending on its phosphorylation status, can thus switch between multiple conformational substates proportional to phosphorylation extents. Because our study relies exclusively on phosphomimetic substitutions, the heterogeneity observed here is likely an underestimate. In contrast to phosphomimetics, phosphorylation introduces two additional negative charges per modification, which may further shift the conformational ensemble toward more unfolded-like states.

**Figure 6.**
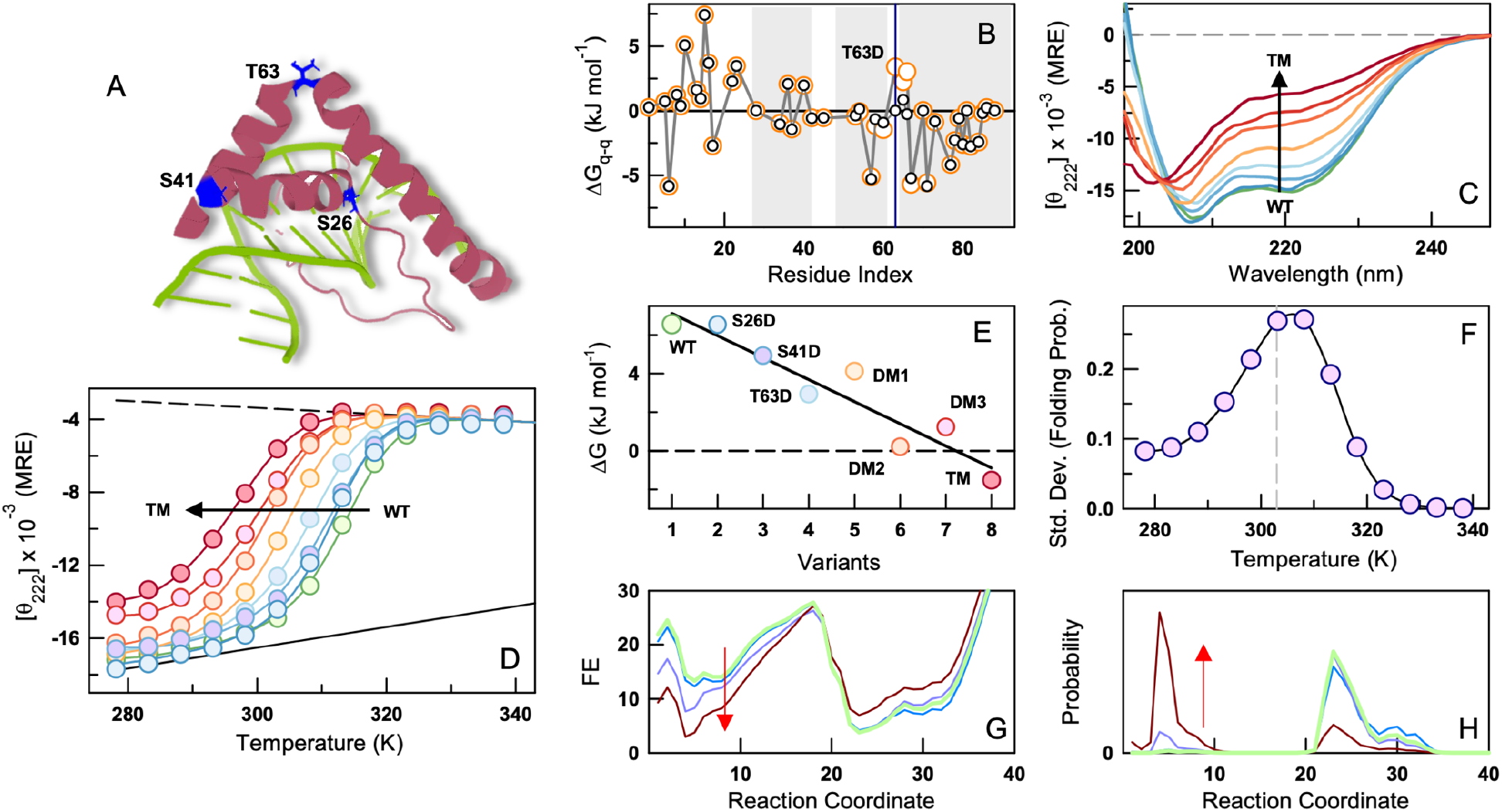
Phosphomimetic mutations destabilize the native ensemble and enhance conformational heterogeneity. (A) Cartoon representation of Nhp6A with DNA (PDB id: 1J5N) highlighting the location of the three mutated sites. (B) Tanford–Kirkwood (TK) electrostatic interaction free-energies as a function of residue number for the WT (black) and a phosphomimetic mutant T63D (orange), respectively. Helices are shaded in gray. The blue vertical line denotes T63 which when mutated to aspartate (i.e. T63D mutant) enhances the electrostatic frustration locally. (C) Far-UV CD spectra of the phosphomimetic mutants at 303 K. The arrow indicates the decrease in helicity. (D) Thermal unfolding curves of the phosphomimetic variants as monitored by far-UV CD at 222 nm (circles). The signals are reported in mean residue ellipticity (MRE) units of deg. cm^2^ dmol^-1^. The curves are a global two-state fit with fixed folded (black line) and unfolded baselines (dashed black line; see text for more details). (E) Thermodynamic stabilities at 298 K. DM1-S26D/S41D, DM2-S41D/T63D, DM3-S26D/T63D, TM-S26D/S41D/T63D. The line through the points is shown to guide the eye. (F) Standard deviation of the folded state probabilities for the different variants at every experimental temperature. The heterogeneity is maximal at ∼303-308 K. The vertical dashed line shows the optimal growth temperature of yeast. (G, H) One-dimensional free energy profiles (panel G) and the corresponding probability densities (panel H) for the WT Nhp6A (green), S26D (blue), T63D (cyan) and TM (dark red) from the bWSME model.

### Stability and DNA binding affinity

To address whether the progressive destabilization of the phosphomimetics correlate with weakened DNA binding, we perform ensemble anisotropy experiments on a few variants. For the mildly destabilizing S26D, we find that the anisotropy of Cy3-labeled DNA displays a rightward shift as a function of S26D concentration relative to the WT (Figure 7A). No discernible transition is observed at concentrations below 130 nM, indicative of a reduced DNA-binding affinity and the absence of the major binding mode. Interestingly, the atomistic MD simulations of this mutant show that the IDR makes fewer contacts with the DNA (Figure S6), likely reducing the interaction strength, and producing a limited bending of the DNA relative to the WT. The anisotropy, a measure of the rotational correlation time, attains the same maximal value as the WT suggesting that the protein-DNA assembly size is comparable to that of the WT. Remarkably, T63D which is the most destabilized single mutant binds to DNA following a near-identical trend as the WT. Again, simulations are consistent with these trends, as indicated by the similar bending induced by the WT and T63D mutant (Figure S6B). The triple mutant (TM) that harbors both these mutations apart from S41D and the most destabilized variant in the set studied, shows a binding isotherm which is shifted farthest to the right and does not exhibit the major binding mode. Taken together, it is apparent that S26D mutation has the maximal effect on the DNA binding as the mutation introduces a negative charge right at the binding interface with DNA, while simultaneously eliminating a H-bond between the hydroxyl group of S26 with A8 N3. Since T63D mutation does not interfere directly with DNA binding, the binding-competent conformation of the mutant is likely selected through a conformational selection mechanism,^71,77^ aided by the strong intermolecular interactions that are preserved. The TM includes both these mutations and thus the DNA complex is even more destabilized.

The fits from the non-specific DNA binding model account for the shape of the isotherm very well (curves in Figure 7A). For T63D, the major and minor binding free energies are comparable to those of the WT, whereas for S26D and TM, the experimental data are best accounted for solely by the minor binding mode free energies (Figure 7B). The predicted macrostate populations for T63D are near-identical as the WT (as the parameters are within experimental error), but the other two variants sample only 0, *m*, and *mm* macrostates (Figure 7C, 7D). Accordingly, the major binding mode (*M*) appears to represent a ‘specific’ interaction that predominates at lower protein concentrations, whereas the minor binding mode (*m*) reflects a ‘non-specific’ interaction detectable only at higher concentrations. Phosphomimetic substitutions thus progressively modulate the accessibility to these binding modes that could be further tuned by protein concentration.

**Figure 7.**
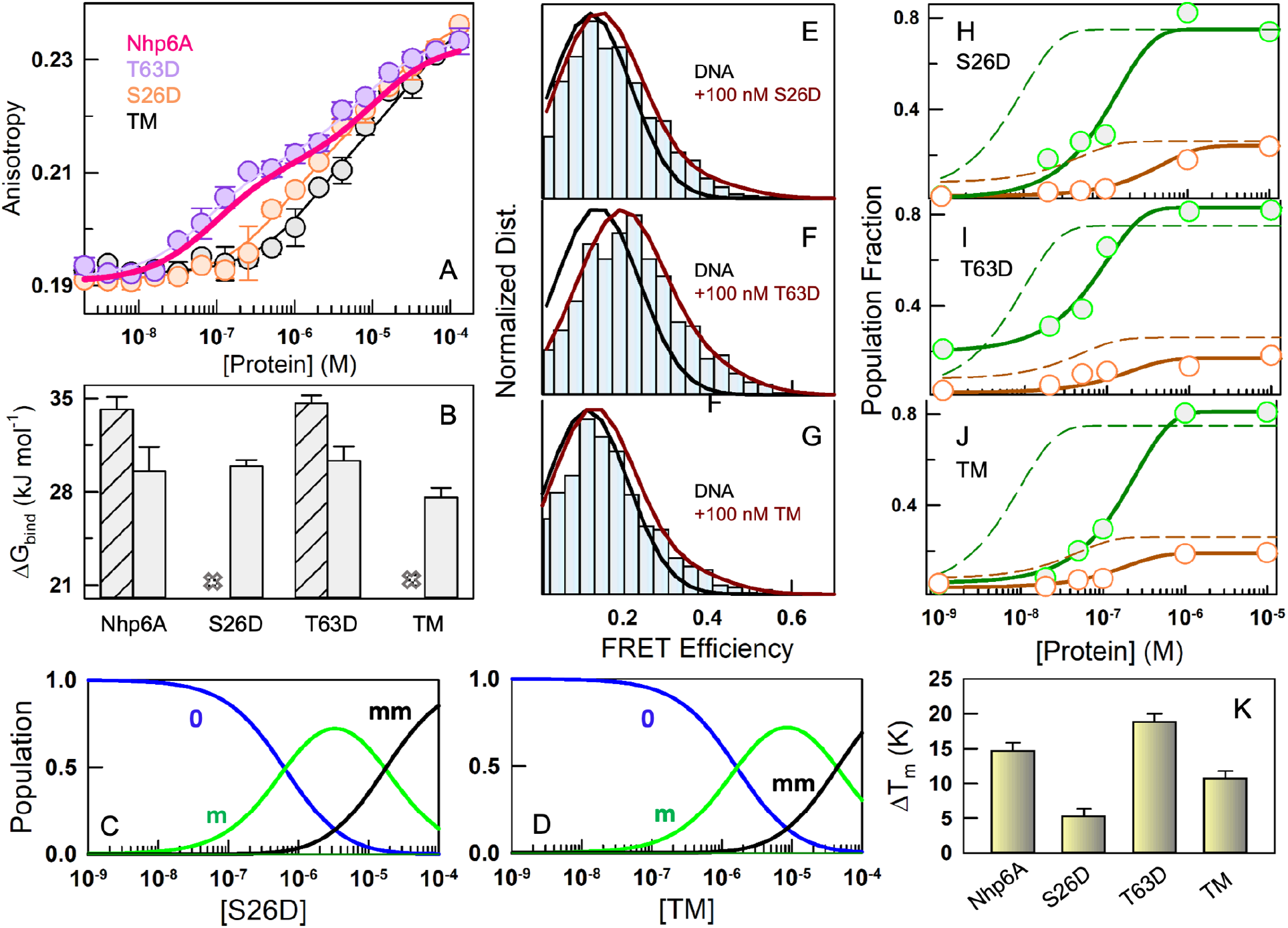
DNA-induced folding and binding characteristics of Nhp6A phosphomimetic variants. (A) Anisotropy of the Cy3-labeled 15 bp dsDNA as a function of protein concentration at 298 K. Note that the T63D variant (purple) displays a similar binding isotherm as that of the WT Nhp6A (magenta). (B) Specific (crossed bar) and non-specific (open bars) binding free energies extracted from the statistical thermodynamic model. The variants S26D and TM lack the specific binding mode, which are indicated by a cross mark. (C, D) Macrostate populations for S26D (panel C) and TM (panel D). The macrostate populations for the T63D variant are near-identical of the WT (see Figure 3G). (E, F, G) Normalized single-molecule FRET efficiency distributions in the absence of protein (black and marked as DNA) and in the presence of 100 nM protein (bars and dark red). Note that neither S26D nor the TM display a significant shift at this concentration indicative of weak binding. (H, I, J) Species fractions of the low (green) and high (orange) FRET efficiency states as a function of protein concentration. The dashed green and dashed dark red curves are the corresponding species fractions for the WT Nhp6A and is shown for comparison. (K) Differences in melting temperatures between the apo (DNA-free) and holo (DNA-bound) forms of the variants from far-UV CD calculated as Δ*T_m_* = *T_m_*_,ℎ*olo*_ − *T_m_*_,*apo*_.

Measures of DNA bending from smFRET experiments follow the trend of ensemble anisotropy measurements for S26D and TM; they induce minimal or no bending at low protein concentrations, as observed from overlapping DNA FRET efficiency histograms in the absence and presence of the protein (Figure 7E-7G, S7). Both low and high FRET species populations reach saturation only at 15-50 fold higher protein concentrations in comparison with the WT (Figure 7H-7J). Since the absolute values of the FRET efficiencies (and by extension the binding angles) do not change, it is possible that more than one molecule binds and bends the DNA which is not captured by smFRET. Surprisingly, T63D displays a smFRET trend similar to the other mutants, while mirroring the WT in ensemble anisotropy experiments. Indeed, the T63D variant is highly stabilized in the holo form relative to S26D and TM (Figure 7K, S8); this observation demonstrates that the binding modes of T63D and WT lead to similar overall thermodynamic stabilization, and by extension a similar binding mode. Taken together with the statistical modelling and molecular simulation results (Figure S6), we conclude that WT and T63D bind in the major binding mode (*M*) at lower concentrations, while S26D and TM bind only via the minor binding mode (*m*).

### Conservation of stability-folding signatures across Nhp6A orthologs

Finally, we ask whether these mechanisms and observations hold across Nhp6A orthologs. A BLAST search within the *Saccharomyces cerevisiae* strains reveals a near perfect conservation of sequence (Figure S9). On the other hand, substantial sequence variability is observed upon a wider search within the fungal taxa (Methods; Figure 8A). Sequence positions contributing to structural stability as identified in this work –Y28, W59, Y70, Y81, Y88 – are fully conserved with very low or zero sequence entropy, apart from the functional positively charged residues across the ordered region, glycine residues in H2, and binding motifs in the N-terminal IDR including the two proline residues. At position 41, both glutamate and alanine residues (69%) are observed apart from serine (28%), indicating that our observations on S41D need not carry over to orthologs. Serine at position 26 is, however, 100% conserved highlighting the critical interaction with DNA and whose phosphorylation not only abrogates this interaction but enhances the electrostatic repulsion with the phosphate backbone. Position 63 is occupied primarily by threonine (42%), serine (34%), and asparagine (18%).

**Figure 8.**
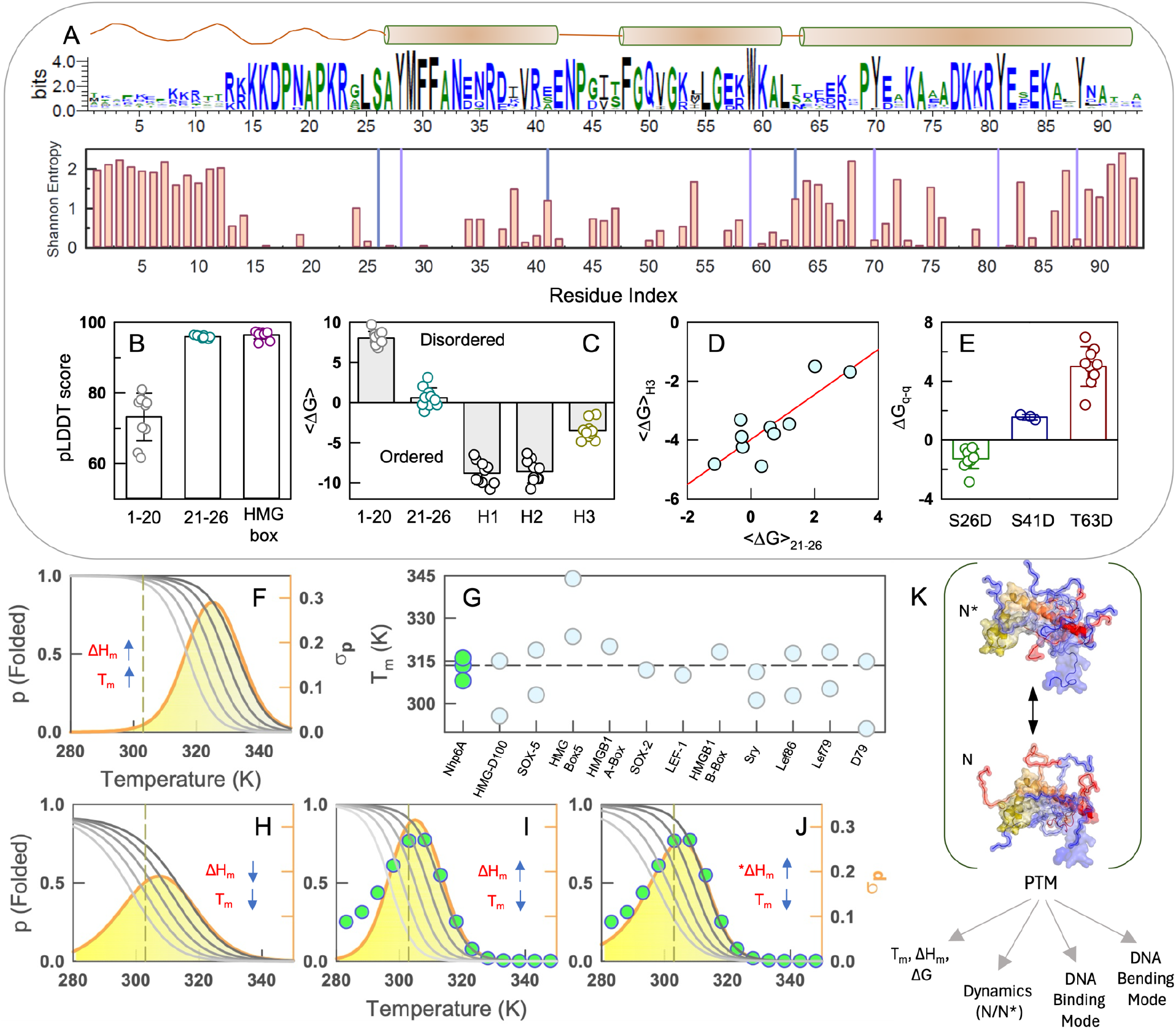
Conserved Nhp6A conformational landscapes, stability determinants and conformational switches. Free energy units are in kJ mol^-1^. (A) Weblogo of diverse sequences from the fungal taxa highlighting strong sequence conservation of different motifs (top; see main text for details). The Shannon entropy gives an informational entropic view of the same (bottom). The vertical purple and blue lines indicate the location of Y/W residues and phosphomimetic mutations, respectively. (B) The predicted local distance difference test (pLDDT) score generated by AlphaFold3 for the 10 selected sequences, grouped into distinct regions. (C) Stability of the different regions as predicted by the bWSME model for the ten chosen orthologs at 298 K. (D) Correlation between the stability of H3 and the motif 21-26. The red line represents the best fit through points with a Pearson’s correlation coefficient of 0.84 and *p*=0.0024. (E) Tanford-Kirkwood electrostatic interaction free energies for aspartate substitutions at positions 26, 41, and 63 for the 10 selected sequences. (F) A simulated scenario for a protein exhibiting high melting temperature (*T*_m_) and enthalpy of unfolding at the midpoint (Δ*H*_m_). The right most unfolding curve belongs to the WT while the subsequent unfolding curves to the left are that of mutants whose melting temperatures are lower (left axis). The orange curve and the associated shaded area plots the standard deviation of folded state probability as a function of temperature (σ_p_; right axis). (G) Melting temperatures of Nhp6A orthologs from literature. Multiple circles for a given protein are indicative of melting temperatures from different structural probes. The dashed horizontal line indicates the melting temperature for Nhp6A from far-UV CD. (H, I, J) Same as panel F but for different assumed values of T_m_ and ΔH_m_. The unfolding curves of the mutants in panel J are simulated assuming a decrease in cooperativity at lower melting temperatures. The circles in panels I and J are the experimentally determined values for Nhp6A and taken from Figure 5F. (K) A cartoon representation summarizing the current work.

We selected 10 sequences spanning phylogenetically diverse fungal taxa (Figure S10) and predicted their three-dimensional structures using AlphaFold3. The predicted structures are highly similar to Nhp6A, with a Cα RMSD of 1.2 Å, and exhibit pLDDT scores exceeding 95 for the ordered helical region, while the N-terminal IDR spanning residues 1-20 scores substantially lower, consistent with its disordered nature (Figure 8B). The region 21-26 that abuts H3 forming a part of the ‘extended strand’ (residues 18-25),^30^ is predicted to be highly ordered. Free energy profiles computed using the bWSME model reveal broad native wells and folding barriers comparable to those of Nhp6A (**Figure S11**). Notably, intermediate-like excited states are populated on the native side of the folding barrier (arrow, Figure S11); these correspond to partially structured conformations in helix 3, as evidenced in the helix-level stability plots (Figure 8C). Helix 3 is consistently the least stable structural element across all orthologs examined, mirroring our observations in Nhp6A. Interestingly, the region 21-26 is neither fully folded nor fully disordered exhibiting local stability values close to zero (Figure 8C). The stability profiles of H3 are highly correlated with that of the residues 21-26 indicating their thermodynamically coupled nature (Figure 8D; also see Figure 2D). To assess whether these signatures extend to phosphomimetic perturbations, we performed Tanford–Kirkwood electrostatic free energy calculations on the selected subset by substituting S26, S41/T41, and S63/T63 with aspartate. In all orthologs, introduction of a negative charge at position 63 produces more unfavorable electrostatic interactions than equivalent substitutions at positions 26 or 41 (Figure 8E), underscoring the critical role of position 63 in governing the structural integrity of Nhp6A orthologs.

From the perspective of a conformational switch, multi-site phosphorylation (or any multi-site post-translational modification (PTM) or perturbation) could tune protein thermodynamic stability (Δ*G*) in a continuous rheostat-like manner, lowering the melting temperatures (*T_m_*) thereby reducing the population of the folded state (*p*). Because Δ*G* and *p* are related through the expression *p* ∝ *e*^−Δ*G*⁄*RT*^, changes in population are significant even for small changes in stability, and this is particularly amplified when the cooperativity of transition is larger (i.e. larger Δ*H_m_*).

However, such stability tuning is functionally relevant only when PTM-induced shifts in *T_m_*s span the physiological growth temperature of an organism. If the melting temperature is high (e.g. 333 K), PTM-driven shifts in stability would not substantially alter the folded-state population at a growth temperature of, say, 303 K — as evident from the standard deviation of the folded-state probability across temperature (σp; orange in Figure 8F and right axis). A comparison of the melting temperatures reported in the literature across eukaryotic Nhp6A orthologs studied by various groups reveals conservation of a low melting temperature (∼314 K) for this fold, independent of its sequence or phylogenetic origin (Figure 8G).^24,70,71,75,78–81^ Under such low-*T_m_* conditions, all three cooperativity scenarios - low cooperativity (low Δ*H_m_*), high cooperativity (high Δ*H_m_*) or mutation-dependent cooperativity (*Δ*H_m_*; mutation reducing the cooperativity at increasing destabilization values) – lead to substantial changes in folded state population at 303 K, the optimal growth temperature of yeast (Figure 8H-8J). Indeed, the best agreement with the experimentally derived σp values (green circles in Figure 8I, 8J) is observed for the two high cooperativity scenarios. Thus, the coupling of low Tm with high Δ*H_m_* (i.e. an unfolding cooperativity comparable to that of larger proteins) in Nhp6A is likely an evolutionarily selected feature, enabling maximal sensitivity to phosphorylation-driven perturbations.

## Discussion

The central finding of this work is that Nhp6A occupies a distinct thermodynamic regime, a folded core stabilized by unusually large unfolding cooperativity and low melting temperature (∼313 K; Figure 1), in addition to the positively charged N-terminal intrinsically disordered region. Despite the large cooperativity of the folding transition, the native ensemble is not a single folded state but comprises at least two subensembles, N and N* (Figure 2). These two states differ primarily in the conformational status of the C-terminal turns of helix 3, which is coupled to the extent of structure formed by residues 21–26 — the region that constitutes part of the N-terminal IDR. A second, distinct source of heterogeneity arises from alternate packing within the folded core, driven by partial melting of helix 2 (Figure 5). This structural plasticity has direct structural consequences for how Nhp6A engages DNA. Ensemble anisotropy resolves two distinct DNA-binding modes with different affinities, while smFRET measurements identify two DNA-bending geometries (Figures 2, 4). Given that the 15-bp DNA duplex used here presents two minor grooves, this structural degeneracy is well-suited to permit close packing of multiple Nhp6A molecules along a DNA lattice — a plausible requirement for its role in global chromatin accessibility, where Nhp6A must bind DNA at high density without steric clashes. Moreover, the weaker binding mode observed at high protein concentrations (>1 µM; Figure 3) falls within the range of Nhp6A concentrations estimated in the yeast nucleus (30–40 µM),^32^ suggesting it could be functionally relevant under these physiological conditions.

Single-molecule DNA diffusion experiments and simulations on Nhp6A by Johnson and co-workers provide strong evidence for at least three diffusion coefficients of Nhp6A diffusing on DNA.^82^ Their coarse-grained simulations further identify two modes of binding: one in which the entire protein is bound to DNA, and another in which only the N-terminal IDR remains bound while the rest of the protein is more loosely associated. These observations are consistent with a requirement for two binding modes in DNA binding proteins — one that facilitates rapid scanning along DNA and another that supports stable, high-affinity binding, with a strong coupling between folding and binding.^83–86^ Taken together with our identification of N and N* within the native ensemble (Figure 2), and the enhancement of N* in the presence of DNA, we hypothesize that N and N* correspond to the same two binding modes computationally identified by Johnson and co-workers.^82^ In this model, the stronger binding mode would map onto N* (the more folded subensemble), while the weaker, more dynamic binding mode would correspond to N (the less folded subensemble).

Phosphorylation introduces another layer onto this complex binding-bending landscape, particularly from the perspective of the two phosphomimetic substitutions S26D and T63D (Figure 6). S26D, abolishes the major DNA-binding mode while leaving global stability largely unaffected — a mutation that acts locally at the binding interface without disrupting the folding equilibrium. On the other hand, T63D substantially destabilizes the folded ensemble through disruption of a favorable local electrostatic environment near the stabilizing H3 N-terminal segment, yet binds DNA with an affinity and macrostate distribution nearly indistinguishable from WT (Figure 7). This is a counterintuitive result under most simple fitness–stability relationships,^87–90^ which generally predict that destabilization should proportionately compromise function. Here, DNA appears to rescue the mutant by overcoming the unfavorable intramolecular interactions in T63D through favorable intermolecular interactions, thus acting as a chaperone. This is not a one-off observation; there is strong evidence for DNA acting as a folding chaperone for the disordered CytR DNA-binding domain,^36,91,92^ the destabilized Y19A mutant of FruR,^93^ and the Brk repressor DBD.^94^ Indeed, the NMR spectra of the LEF1 HMG-protein is suggestive of a disordered protein while it recovers the dispersion expected of folded systems in the presence of DNA.^80^ It thus appears that phosphomimetic substitutions can act at multiple levels, tuning any or all of the following conformational features: the relative population of N and N*, global thermodynamic stability, DNA-binding mode, and the extent of DNA bending (Figure 8K).

The observed decoupling of folding-binding equilibria in T63D raises questions on the functional logic of phosphorylating a site that does not contact DNA. One possibility, consistent with the rheostat-like destabilization we observe across the single, double, and triple mutants, is that such sites (i.e. T63-like positions) are not intended to abolish binding but rather to shift the conformational equilibrium toward more unfolded-like states — states that may be preferentially recognized by cellular quality-control or degradation machinery. In this model, T63 phosphorylation would function less as an on-off binding switch and more as a rheostat controlling the population of (un)folded Nhp6A. However, S26 phosphorylation will directly act as a binding-specific switch. The near-perfect conservation of S26 across fungal taxa, contrasted with substantial polymorphism at position 63 (Thr/Ser/Asn), is consistent with our expectation: the binding-critical residue is invariant, while the stability-modulating residue tolerates chemically conservative substitutions (Figure 8).

There are, however, a few caveats. First, despite the coarse agreement between experiments, MD simulations and the statistical model, the extent of structural heterogeneity in the apo form requires to be fully ascertained. This will form the blueprint for mapping the holo-Nhp6A landscape at a quantitative level, the sampling of which is hampered by the rough landscape of the N-terminal IDR. While NMR experiments on Sox2 HMG protein do point to the possibility of multiple partially structured states in the apo form,^71^ more detailed mutagenesis-based experiments are required to discern them experimentally in Nhp6A. Likewise, the DNA-bending angle recovered from atomistic MD is lower than those measured by smFRET, most likely reflecting the restraining potential used to preserve Watson-Crick pairing during enhanced sampling or a mismatch between the angle definitions between the smFRET and the atomistic MD; this limits our ability to assign the two bending states to specific atomistic geometries with full confidence, and fully unrestrained simulations will be needed to resolve this gap. Second, phosphomimetic substitutions introduce a single negative charge and cannot fully recapitulate the doubly negative phosphate group; our results should therefore be regarded as a lower bound on the conformational heterogeneity accessible to phosphorylated Nhp6A. Finally, through a combination of experimental and computational approaches we capture the direction and relative magnitude of PTM-induced perturbations well, but is not intended as a quantitative predictor. In-cell studies measuring the stability and DNA binding of Nhp6A and its phosphorylated variants, though challenging, could provide a more nuanced view of the interplay of the various thermodynamic factors in the cellular context.

Taken together, Nhp6A exemplifies a design principle wherein marginal stability, large unfolding cooperativity, and electrostatic frustration at evolutionarily conserved sites combine to imprint charge-dependent conformational switching on the folding landscape. Our simulations of alternative stability-cooperativity scenarios show that the combination of these features best reproduces the experimentally observed temperature-dependence of heterogeneity in folded state probability (Figure 8), with the maximal heterogeneity coinciding with the optimal growth temperature of *S. cerevisiae*. Marginal stability is thus necessary but not sufficient and it is the pairing with high cooperativity that converts Nhp6A into a molecular switch. Because the ordered HMG-box fold and the disordered N-terminal tail are shared across the wider HMGB protein family — including double-box and acidic-tail-bearing members already known to be regulated by phosphorylation — the sequence-ensemble-dynamics code identified here for Nhp6A may be a template for how architectural, non-sequence-specific DNA-binding proteins couple post-translational signaling to chromatin accessibility.

## Author Information

Corresponding Authors Tel: +91-44-2257 4140

## Abbreviations

HMG: high mobility group
DSC: differential scanning calorimetry
CD: circular dichroism
DBD: DNA binding domain
MD: molecular dynamics
smFRET: single molecule Förster resonance energy transfer
WSME: Wako-Saitô-Muñoz-Eaton

## Acknowledgements

A.N.N. acknowledge the FIST facility sponsored by the Department of Science and Technology (DST, India) at the Department of Biotechnology, IIT Madras (Chennai, India) for the instrumentation. A.N.N. and S. T. are grateful for support from the India Japan Science and Technology Cooperation Programme grant no. DST/INT/JSPS/P-394/2024 (G). Y.I. is grateful for JSPS KAKENHI Grants JP24K18075. D.D.S. receives support from the Basque Government through Project IT2067-26 and grant PID2024-158678NB-I00 funded by MICIU/AEI/10.13039/501100011033 and “ERDF A way of making Europe”.

## Data Availability Statement

The data underlying this article are available in the article and in the online supplementary material.

