## Supporting Information for "Switching Functional DNA-Binding Modes by Tuning Protein Order-Disorder Equilibria"

AUTHOR INFORMATION

**Corresponding Authors**

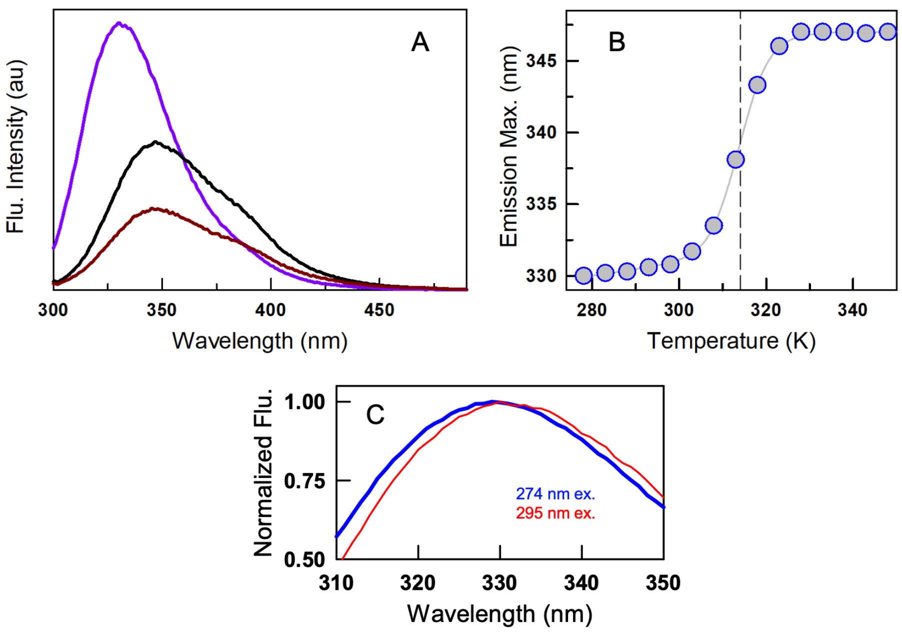


**Figure S1** (A) Fluorescence emission spectra of Nhp6A at 298 K (purple), 333 K (black) and 358 K (dark red) upon excitation at 295 nm. (B) The wavelength of maximum fluorescence emission as a function of temperature indicates a two-state-like transition with a melting temperature (vertical dashed line) of 314.5 K. (C) Fluorescence emission spectra of Nhp6A at the excitation wavelengths indicated.

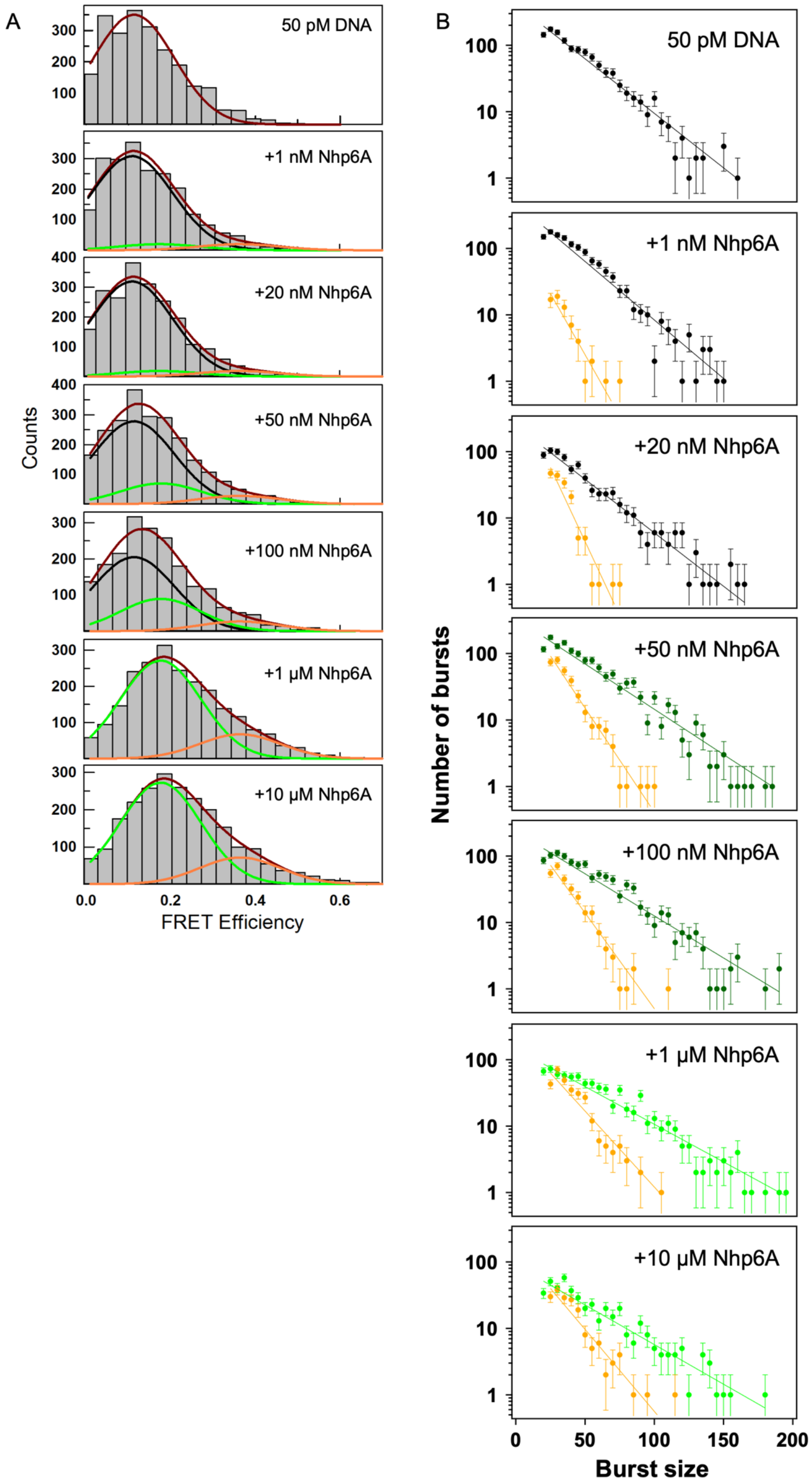


**Figure S2** (A) Same as a Figure 4A in the main text, but for all the protein concentrations explored. (B) Similar to a Figure 4C in the main text, but for all the protein concentrations explored. Based on the predominant component of detected bursts with their FRET efficiency $E$ less than 0.20, they are shown in black (DNA), dark green (DNA and low FRET state) and green (low FRET state), respectively.

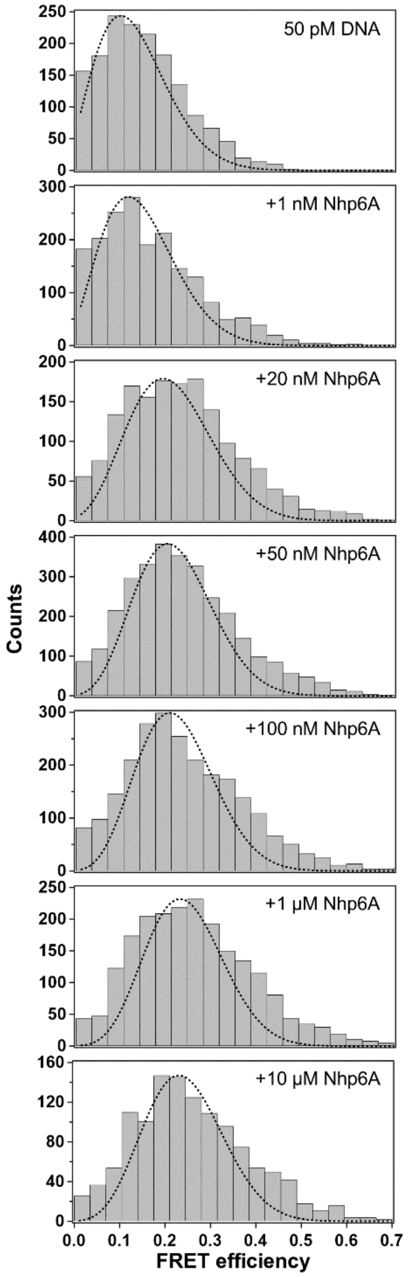


**Figure S3** smFRET data and the expected shot noise width estimated by assuming a single conformation and averaged photon counts. The theoretical shot noise widths were estimated by assuming that all bursts had the same donor and acceptor photon counts as well as the background counts determined by averaging the experimental counts under respective conditions. It should be noted that the estimated widths should be considered the lower limit of the shot noise widths since the photon numbers of the actual bursts were not constant. The calculation was performed using a homemade Python program based on eq. S7 in Supporting Information of a previous report.[1]

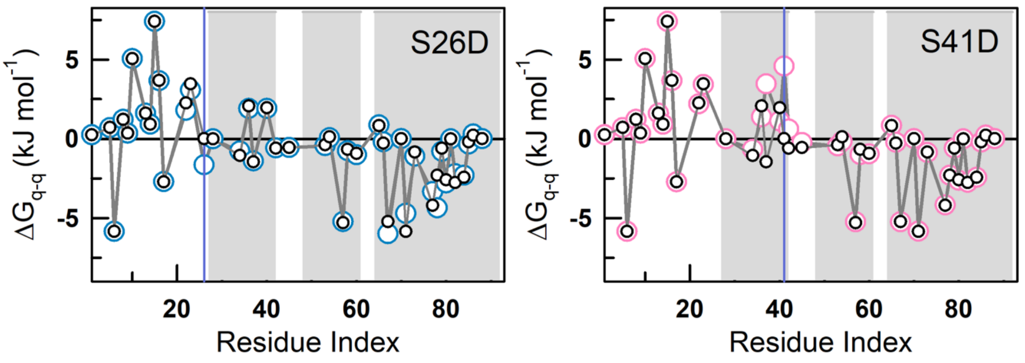


**Figure S4** Tanford-Kirkwood analysis of S26D and S41D variants (colored circles) relative to the WT Nhp6A (black circles). The vertical lines indicate the mutated position.

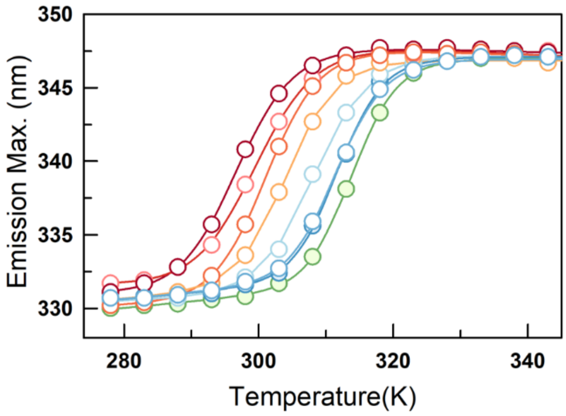


**Figure S5** Emission maximum of W59 as a function of temperature for the mutants studied in this work, following the color code in Figure 5E (main text).

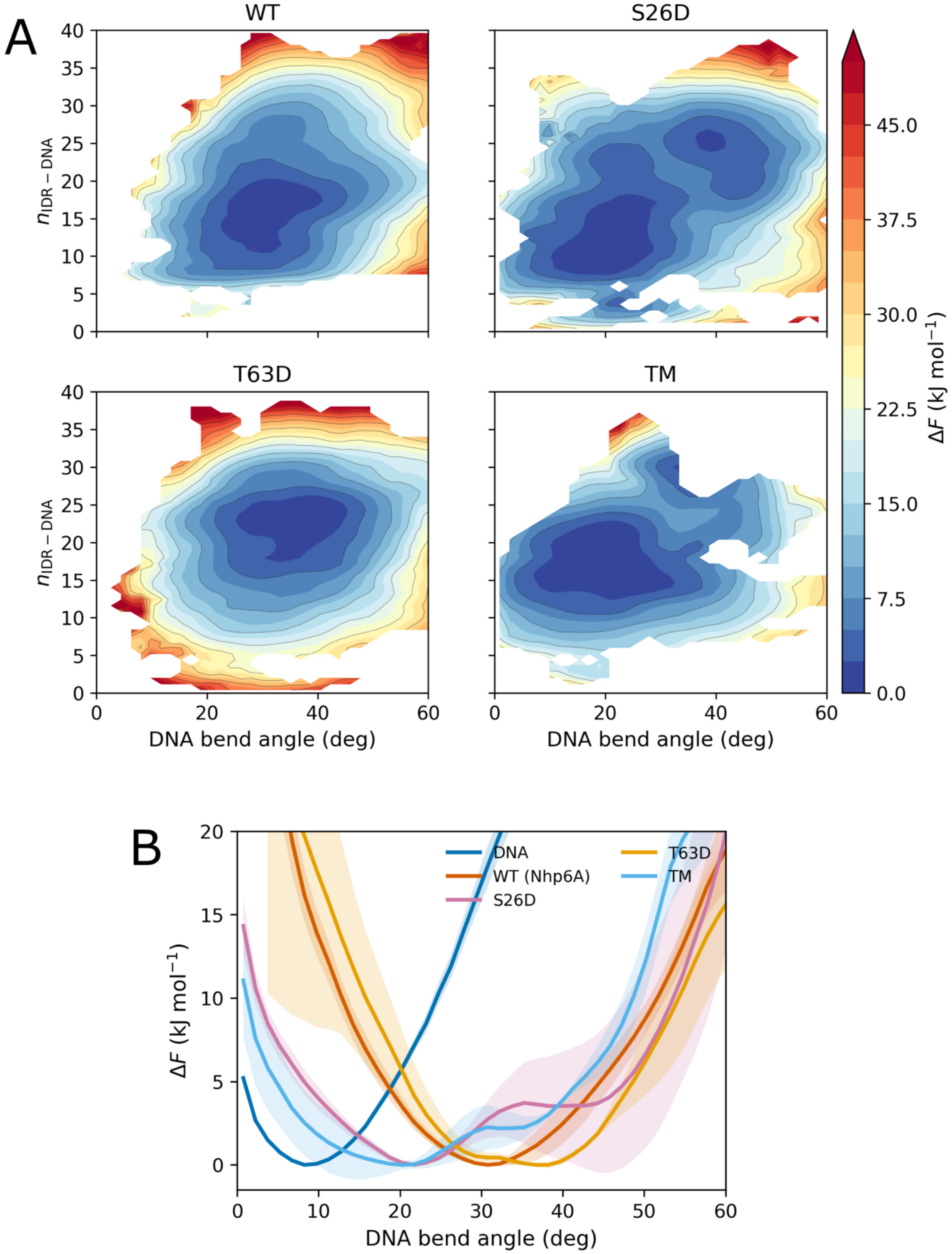


**Figure S6** (A) Free energy landscape for the protein:DNA complex as a function of the bending angle and the number of IDR-DNA contacts (nIDR-DNA) for the WT and selected mutants. (B) Potential of mean force for the projection on the bending angle of the DNA for the different simulation datasets.

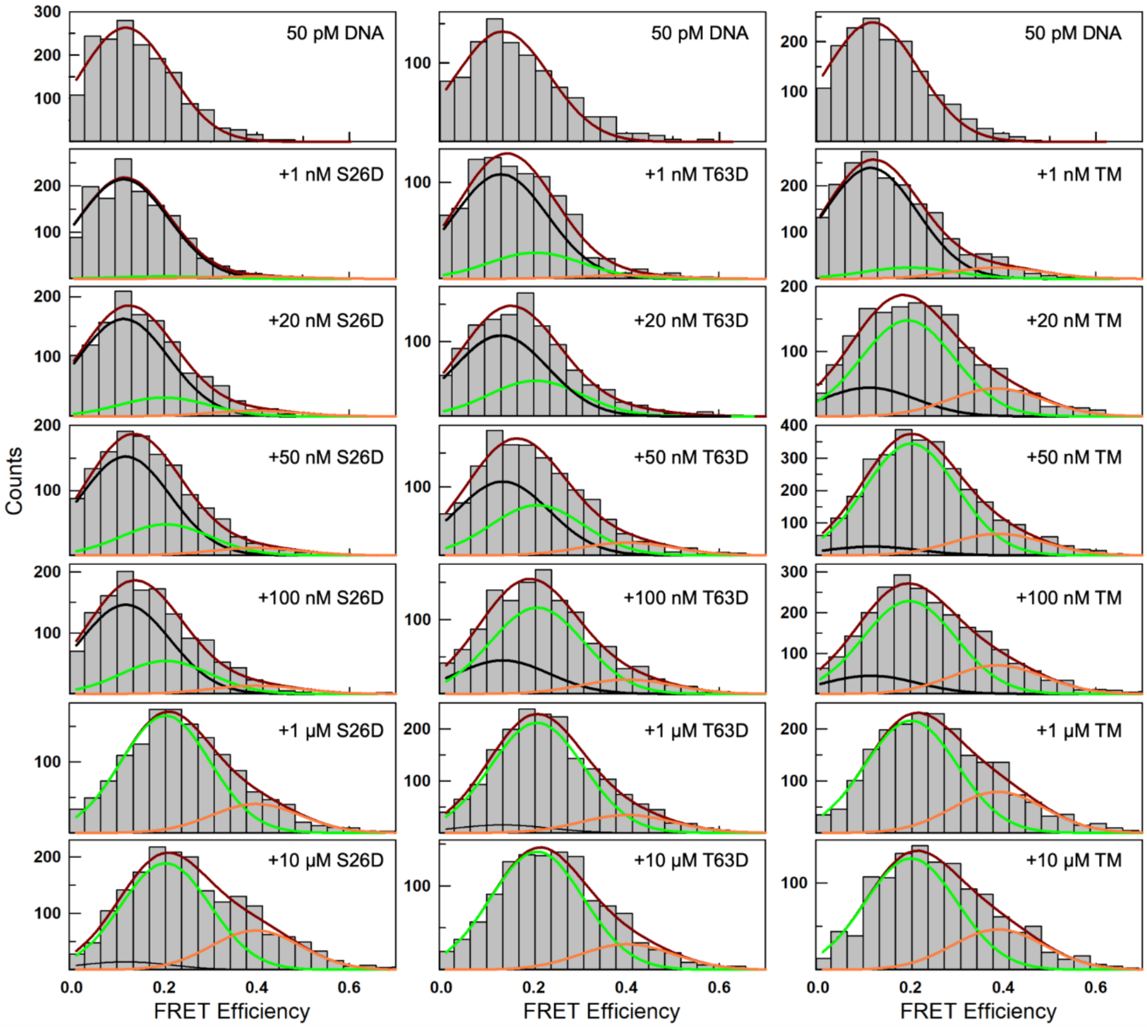


**Figure S7** Same as a Figure S2, but at all the protein concentrations explored for S26D (left column), T63D (middle column) and TM (right column). Note that the y-axis scale varies across plots to aid visualization.

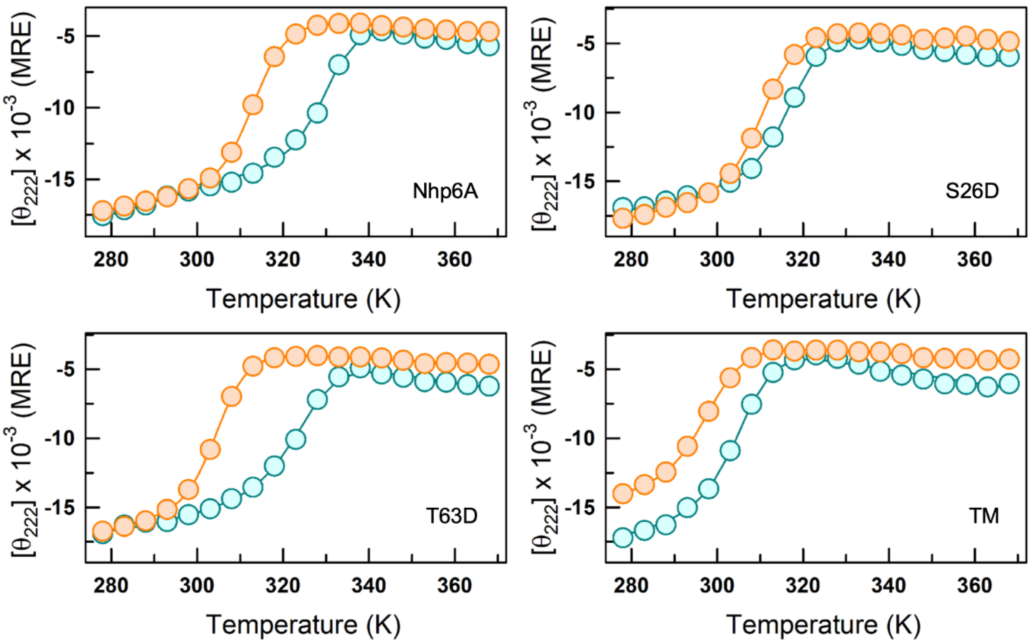


**Figure S8** Far-UV CD signal at 222 nm as a function of temperature for the apo (orange) and holo (DNA-bound; cyan) forms of the variants indicated.

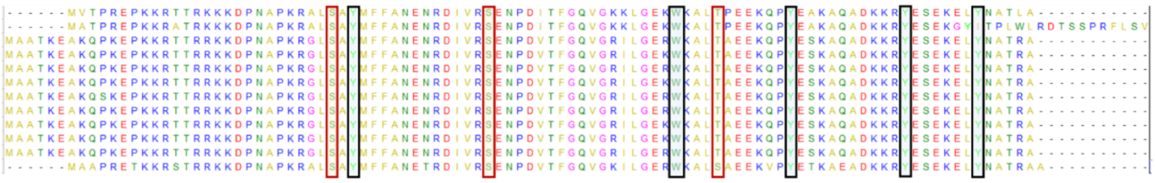

**Figure S9** Sequence alignment of Nhp6A from *Saccharomyces cerevisiae* strains. Residues of interest in this study (highlighted) are fully conserved.

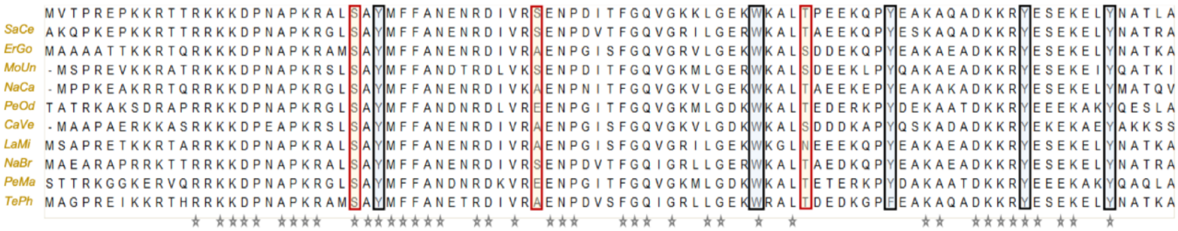


**Figure S10** Sequence alignment of Nhp6A orthologs from different fungi in the following order (the first sequence is Nhp6A from *Saccharomyces cerevisiae*): *Saccharomyces cerevisiae* (SaCe), *Eremothecium gossypii* (ErGo), *Monosporozyma unispora* (MoUn), *Naumovozyma castellii* (NaCa), *Penicillium odoratum* (PeOd), *Candida verbasci* (CaVe), *Lachancea mirantina* (LaMi), *Nakaseomyces bracarensis* (NaBr), *Penicillium manginii* (PeMa), *Tetrapisispora phaffii* (TePh). Stars mark the positions that are invariant across all sequences.

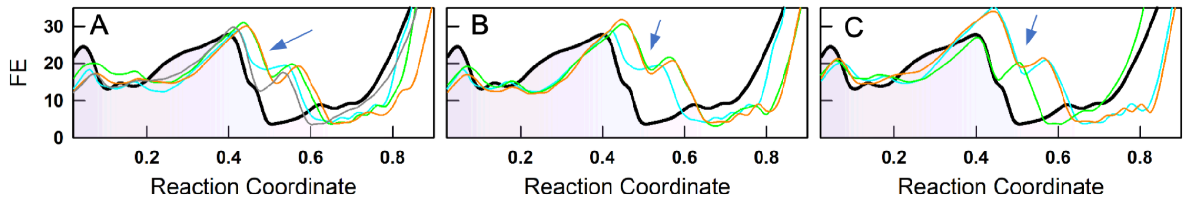


**Figure S11** Free energy profiles of Nhp6A orthologs predicted by the bWSME model at 298 K, relative to the Nhp6A from *Saccharomyces cerevisiae* (black), grouped into three panels for ease of viewing. The color coding is as follows: *Saccharomyces cerevisiae* (SaCe; Nhp6B, cyan in panel A), *Eremothecium gossypii* (ErGo; green in panel A), *Monosporozyma unispora* (MoUn; orange in panel A), *Naumovozyma castellii* (NaCa ; dark gray in panel A), *Penicillium odoratum* (PeOd; cyan in panel B), *Candida verbasci* (CaVe; green in panel B), *Lachancea mirantina* (LaMi; orange in panel B), *Nakaseomyces bracarensis* (NaBr; cyan in panel C), *Penicillium manginii* (PeMa; green in panel C), *Tetrapisispora phaffii* (TePh; orange in panel C). Arrow marks in the position of the intermediate characterized by significant unfolding at the C-terminal of H3.

**Table S1** Simulation datasets used in this study.

| Systems | Type of run | Runs/Walkers | Length/Walker (ns) | Aggregate (ns) |
|---|---|---|---|---|
| DNA | Equilibrium | 3 | 250 | 750 |
| DNA | Metadynamics | 6 | 100 | 600 |
| Nhp6A Apo | Equilibrium | 3 | 250 | 750 |
| Nhp6A Apo | Metadynamics | 6 | 1250 | 7500 |
| Nhp6A:DNA | Equilibrium | 3 | 250 | 750 |
| Nhp6A:DNA | Metadynamics | 6 | 350 | 2100 |
| Nhp6$A_2$:DNA | Equilibrium | 3 | 250 | 750 |
| S26D:DNA | Metadynamics | 6 | 350 | 2100 |
| T63D:DNA | Metadynamics | 6 | 350 | 2100 |
| TM:DNA | Metadynamics | 6 | 300 | 1800 |

**Table S2** Parameters derived from a global two-state fit to the far-UV CD monitored thermal unfolding curves of the variants.

| **Protein** | $T_m$ **(K)** | $\Delta H_m$ **(kJ mol$^{-1}$)** |
|---|---|---|
| WT | 313.4 ± 0.5 | 185.9 ± 14.1 |
| S26D | 311.4 ± 0.5 | 197.4 ± 17.4 |
| S41D | 310.7 ± 0.5 | 164.5 ± 10.5 |
| T63D | 304.4 ± 0.5 | 163.7 ± 10.1 |
| DM1 | 307.9 ± 0.5 | 163.2 ± 10.0 |
| DM2 | 298.7 ± 0.5 | 116.1 ± 6.0 |
| DM3 | 300.8 ± 0.4 | 147.8 ± 8.1 |
| TM | 294.1 ± 0.5 | 99.2 ± 5.4 |
